# A complex of distal appendage-associated kinases linked to human disease regulates ciliary trafficking and stability

**DOI:** 10.1101/2020.08.21.261560

**Authors:** Abdelhalim Loukil, Chloe Barrington, Sarah C. Goetz

## Abstract

Cilia biogenesis is a complex, multi-step process involving the coordination of multiple cellular trafficking pathways. Despite the importance of ciliogenesis in mediating the cellular response to cues from the microenvironment, we have only a limited understanding of the regulation of cilium assembly. We previously identified a kinase that acts as a key regulator of ciliogenesis, TTBK2. Here, using CRISPR kinome screening, we identify the CK2 subunit CSNK2A1 as an important modulator of TTBK2 function in cilia trafficking. Super-resolution microscopy reveals that CSNK2A1 is a centrosomal protein concentrated at the mother centriole and associated with the distal appendages where it physically interacts with TTBK2. Further, *Csnk2a1* knockout partially corrects defects in cilia formation and length in *Ttbk2* hypomorphic cells. *Csnk2a1* mutant cilia are longer than those of control cells and exhibit instability, particularly at the tip. *Csnk2a1* mutant cilia also abnormally accumulate key cilia assembly and SHH-related proteins including IFT, GLI2, KIF7, and Smoothened (SMO). *De novo* mutations of *Csnk2a1* were recently linked to the human genetic disorder Okur-Chung neurodevelopmental syndrome (OCNDS). Consistent with the role of CSNK2A1 in cilium stability, we find that expression of OCNDS-associated *Csnk2a1* variants in wild-type cells cause ciliary structural defects. Our findings provide new insights into mechanisms involved in ciliary length regulation, trafficking, and stability that in turn shed light on the significance and implications of cilia instability in human disease.

**SIGNIFICANCE STATEMENT:** Primary cilia (PC) are sensory organelles that play essential roles during development and adulthood. Abnormal functioning of PC causes human disorders called ciliopathies. Hence, a thorough understanding of the molecular regulation of PC is critical. Our findings highlight CSNK2A1 as a novel modulator of cilia trafficking and stability, tightly related to TTBK2 function. Enriched at the centrosome, CSNK2A1 prevents abnormal accumulation of key ciliary proteins, instability at the tip, and aberrant activation of the Sonic Hedgehog pathway. Further, we establish that *Csnk2a1* mutations associated with Okur-Chung neurodevelopmental disorder (OCNDS) alter cilia morphology. Thus, we report a potential linkage between CSNK2A1 ciliary function and OCNDS.

## INTRODUCTION

Primary cilia are microtubule-based cellular projections that function as important cellular signaling organelles. These structures are required to transduce and integrate signaling through a variety of important developmental pathways, with dysfunction of primary cilia linked to several recessive genetic syndromes. In addition to their roles in embryonic development, cilia also persist on the cells of most tissues during adulthood (1–4).

Cilia are dynamic organelles, and trafficking of proteins and other materials in and out of cilia by the process of intraflagellar transport (IFT) is vital both for the maintenance of these structures as well as for the process of signaling (5–7). In addition, cilia are dynamically assembled and disassembled in both developing embryos and cultured cells (3). Cilia play important roles in multiple cellular signaling pathways that are activated by extracellular signals. It is therefore critical to define the processes and pathways that function upstream of IFT to control the presence or absence of a cilium. In spite of this, our knowledge of such pathways remains limited.

In prior work, we identified a kinase, Tau tubulin kinase 2 (TTBK2), that is required for cilium assembly. TTBK2 mediates both the recruitment of IFT proteins to the mother centriole as well as the removal of centrosomal proteins that suppress cilium assembly (8). In cultured fibroblasts, TTBK2 dynamically localizes to the mother centriole prior to the initiation of cilium assembly. In mouse embryos lacking TTBK2, cilium assembly completely fails, resulting in embryonic lethality. This implies that TTBK2 acts in a pathway that is critical for the regulation of ciliogenesis. TTBK2 is important for maintaining the stability of the ciliary axoneme (9), and both of these requirements for this kinase are linked to a role for TTBK2 in a human neurodegenerative disease (4). A limited number of interactors and effectors of TTBK2 in ciliogenesis have been identified, including the distal appendage proteins CEP164 (10, 11) and CEP83 (12), and a suppressor of ciliogenesis, M-phase phosphoprotein 9 (13). However, the pathway through which TTBK2 functions to mediate cilium assembly and stability is largely undefined.

Here, to uncover additional players in this pathway, we undertook distinct unbiased approaches to identify both new physical and functional interactors with TTBK2. These included using proximity-dependent biotin identification (BioID) to identify TTBK2-proximate proteins, and a CRISPR-based screen to uncover genetic modifiers of a hypomorphic *Ttbk2* phenotype (9). Among the hits identified in these screens, Casein kinase II subunit alpha (CSNK2A1), the main catalytic subunit of Casein kinase II (CK2) (14), came out in both approaches, making it an attractive candidate as an effector of TTBK2 in cilium formation.

CK2 is a widely expressed serine-threonine kinase that is involved in many cellular processes, including cell cycle progression, circadian clock regulation, and WNT signaling (15–17, 17–19). Moreover, mutations in the CSNK2A1 subunit are associated with Okur-Chung syndrome, a dominantly inherited neurodevelopmental disorder that is also characterized by dysmorphic facial features (20). Here, we show that CSNK2A1 acts in opposition to TTBK2 in establishing and maintaining ciliary structure, with knockdown of *Csnk2a1* partially rescuing ciliary phenotypes of *Ttbk2* hypomorphic mutant cells. We also demonstrate that CSNK2A1 localizes to the centrioles and is required for cilia structure and stability. *Csnk2a1* knockout cells have long cilia that exhibit a variety of trafficking defects. Additionally, over-expression of *Csnk2a1* variants associated with Okur-Chung neurodevelopmental syndrome causes ciliary structural abnormalities. Thus, we reveal a novel role for Casein kinase II in cilia regulation and begin to define a kinase network that regulates the stability of these critical signaling organelles, and point to links between ciliary instability and human disease.

## RESULTS

### CSNK2A1 is an effector of TTBK2 at the mother centriole

In order to identify additional components of the TTBK2 dependent pathways that regulate ciliary assembly and/or stability, we initially undertook two distinct unbiased screens: A BioID proximity labeling screen to find proteins that physically interact with TTBK2 and might, therefore, be effectors or upstream regulators of this crucial kinase (**Fig 1A**); and a CRISPR loss-of-function screen to identify proteins that functionally interact with TTBK2 (**Fig 1B**).

**Figure 1:**
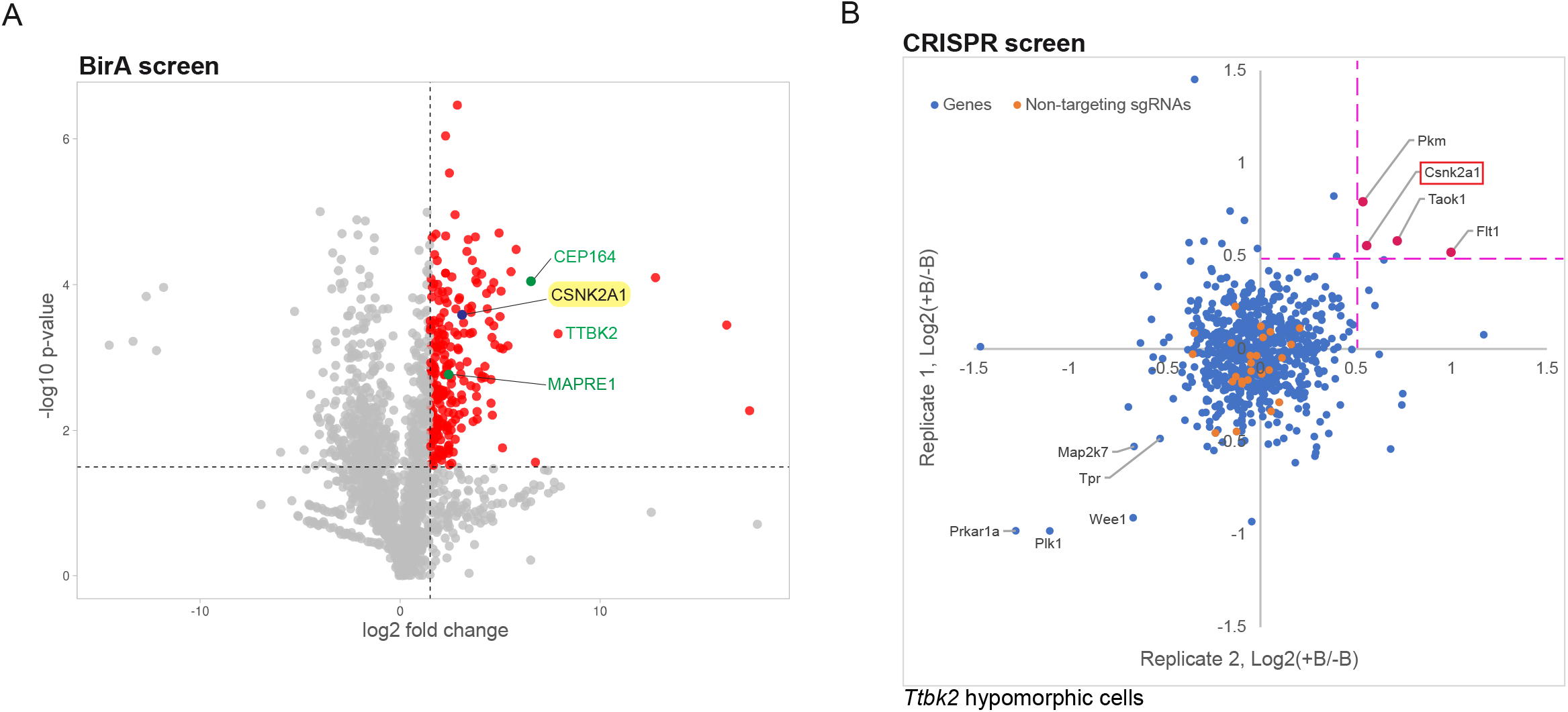
Identification of CSNK2A1 as a putative TTBK2-interacting protein. A. Volcano plot of TTBK2-proximate proteins labeled by BioID. CEP164, TTBK2, and MAPRE1, known interactors of TTBK2, are shown in green. CSNK2A1, a candidate TTBK2-proximate protein is highlighted in yellow. The Y-axis denotes the fold change, which is the log2 of TTBK2: GFP ratio, calculated for each protein. In the X-axis the significance displays the negative log10-transformed p-value for each protein. The red dots show the proteins considered as hits, delimited by the selected thresholds ≥ 1.5 for the log2 fold change and statistical significance. B. The CRISPR screen plot for *Ttbk2* hypomorphic cells. The suppressor screen identifies potential negative modifiers of TTBK2, including CSNK2A1 (red). Each gene is plotted using the mean of the medians extracted from each replicate. Orange dots represent the non-targeting sgRNAs.

In our proximity labeling screen, we tagged TTBK2 at the N terminus with the promiscuous biotin ligase BirA, mutated to be constitutively active (Roux et al., 2012). After verifying that BirA* tagged TTBK2 is active and able to perform its normal cellular function in mediating cilium assembly (**Fig. S1A**), we used the TRex Flp-In system (21) to inducibly express TTBK2-BirA* in HEK-293T cells upon administration of tetracycline. Following induction of the fusion protein and addition of exogenous biotin, cells were lysed, and affinity purification was performed for biotinylated proteins using streptavidin, followed by mass spectrometry (**Fig. S1B, C**). Using this approach, we identified 370 proteins that were enriched by 2-fold or more in cells expressing TTBK2-BirA* compared with the GFP-BirA* negative control (**Fig 1A**, **Supp. Table 1**). Of these 351 were statistically significant at a false discovery rate of 0.01.

In parallel with this approach, we performed a CRISPR loss-of-function kinome screen to identify additional kinases that cooperate with TTBK2 in cilium assembly. We employed a recently described Hedgehog (HH) pathway-responsive reporter, in which 8X GLI response elements drive expression of blasticidin resistance genes (22), in combination with a mouse kinome library (23). We expressed the reporter as well as the kinome sgRNA library (#75316, Addgene; (23) in WT mouse embryonic fibroblasts (MEFs), as well as MEFs, derived from hypomorphic *Ttbk2* mutant embryos (*Ttbk2*^*null/gt*^ trans-heterozygous embryos (9) (**Fig. S2A**). We defined a hit in our screen as any sgRNAs that became enriched within the population following blasticidin selection in the *Ttbk2*^*null/gt*^ cells, but not in the WT MEFs. In other words, hits were sgRNAs that specifically improve the responsiveness of cells to HH pathway activation when TTBK2 activity is impaired. This approach would predominantly identify negative regulators of TTBK2 function either in cilium initiation or cilium stability and maintenance or both (**Fig 1B**, **Fig. S2B Supp. Table 2**). A single kinase emerged as a hit from both of these screens-Casein kinase II subunit alpha (CSNK2A1), the catalytic subunit of Casein kinase II (**Fig 1A, B**). To examine the relationship between CSNK2A1 and TTBK2, we verified that these two kinases associate during ciliogenesis. We co-expressed TTBK2-V5 with GFP or CSNK2A1-GFP in serum-starved HEK 293T cells. We detected an interaction between the two fusion proteins, whereas no interaction was observed in the control pulldown (**Fig 2A**).

**Figure 2:**
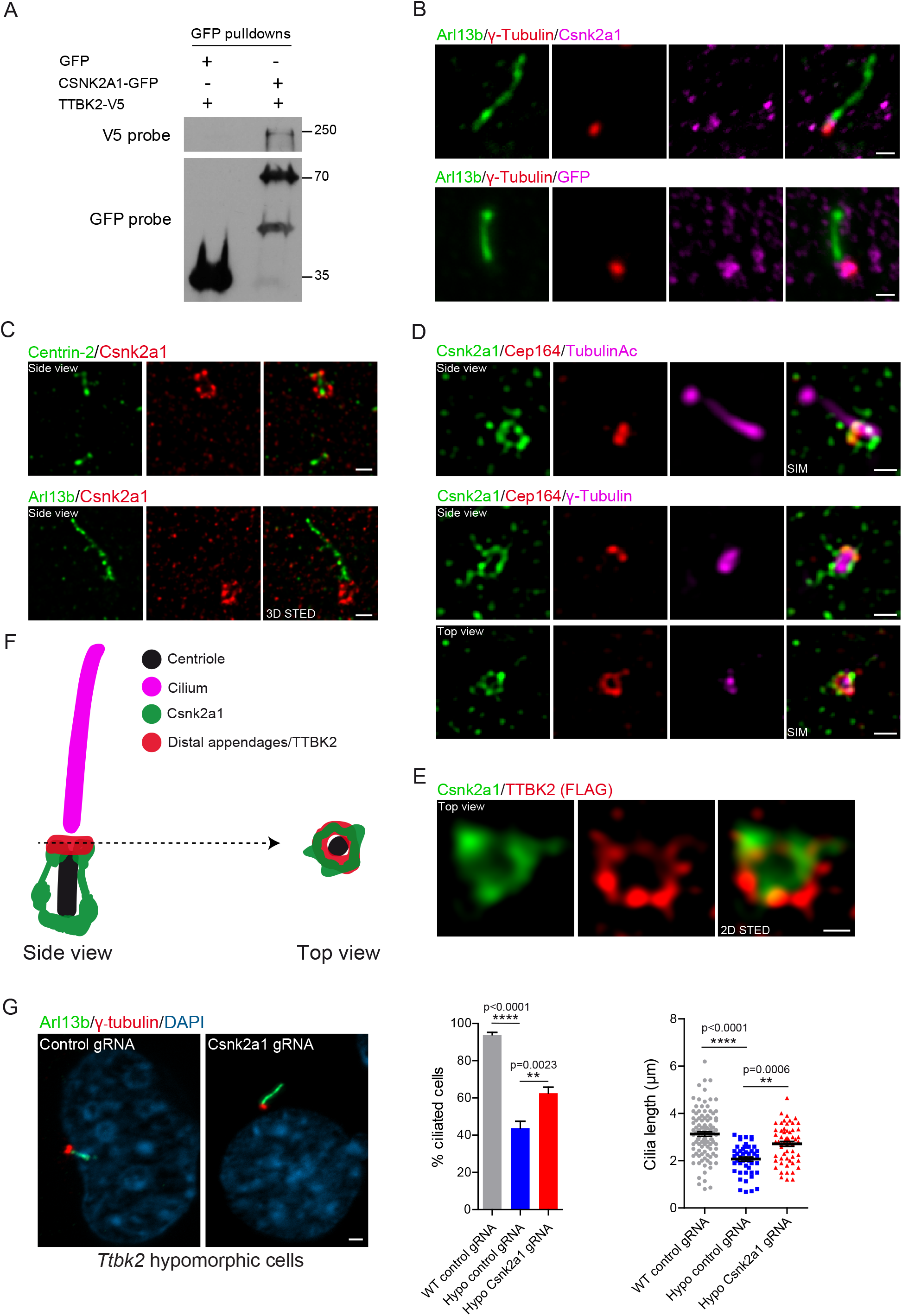
CSNK2A1 is a centrosomal protein that opposes TTBK2 function. A. CSNK2A1 physically interacts with TTBK2 by co-immunoprecipitation. TTBK2-V5 was overexpressed with GFP or CSNK2A1-GFP in HEK293T cells. GFP pulldowns were performed on total cell lysates, and western blots were probed with antibodies to V5 and GFP tags. B. Endogenous (upper panel) and GFP-tagged (lower panel, stably expressed, both magenta) CSNK2A1 colocalizes with the centrosomal protein γ-tubulin (red) and the most proximal part of the cilium, labeled with ARL13B (green) in WT MEFs. Scale bars: 1 μm C. 3D STED super-resolution images show the endogenous CSNK2A1 (red) predominantly localized at the mother centriole in WT MEFs. CSNK2A1 surrounds the centriole core labeled by Centrin-2 (green, upper panel). The lower panel shows CSNK2A1 signal at the vicinity of the cilium labeled by ARL13B. Scale bars: 500 nm. D. CSNK2A1 partially colocalizes with the distal appendage protein CEP164. SIM acquisitions show that CSNK2A1 exhibits a complex basal body-associated organization. Seen in the upper panel, a side view of CSNK2A1 (green) at the base of the cilium labeled with Acetylated alpha-tubulin (magenta), and partially overlapping with CEP164 (red). CSNK2A1 tightly surrounds the pericentriolar material, labeled by γ-Tubulin, partially overlapping with CEP164 (upper and lower panels, side views). CSNK2A1 forms a ring-like structure colocalizing with the CEP164 ring (lower panel, top view). Scale bars: 500 nm. E. Representative 2D STED images from WT MEFs, exhibiting the top view of CSNK2A1 (green), in a ring-shaped structure and colocalizing with TTBK2 (red). Scale bar: 250 nm. F. Cartoon illustrating the subcellular organization of CSNK2A1 at the basal body. CSNK2A1 displays a symmetrical barrel-like structure, which is distally topped with a ring-like shape, colocalizing with distal appendage proteins. G. Cilia defects in *Ttbk2* hypomorphic cells (*Ttbk2*^*null/gt*^) are partially rescued by *Csnk2a1* depletion. Clonal cell lines were generated with control or Csnk2a1 sgRNAs in *Ttbk2*^*null/gt*^ MEFs. Representative immunofluorescence images of serum-starved cells are shown, labeled with the ciliary marker ARL13B (Green) and the centrosomal protein γ-tubulin (red). DNA was stained with DAPI. Scale bar: 2 μm Graphs show the mean percentage of ciliated cells (Left graph; WT control gRNA n=105; Hypo control gRNA n=251; Hypo *Csnk2a1* gRNA n=282 cells) and cilia length (Right graph; WT control gRNA n=105; Hypo control gRNA n=48; Hypo *Csnk2a1* gRNA n=58 cells). Error bars denote the SEM. Statistical comparison was performed by one-way analysis ANOVA with Tukey’s multiple comparisons test. P-values are shown on the graph (ns: not significant).

TTBK2 localizes to the distal appendages of the mother centriole prior to the initiation of cilium assembly (8, 12, 24). To assess whether CSNK2A1 co-localizes with TTBK2 in that compartment and to resolve the precise sub-ciliary localization of CSNK2A1, we performed conventional as well as super-resolution immunofluorescence microscopy approaches using antibodies against endogenous CSNK2A1 as well as a GFP fusion construct. With both approaches, we found that CSNK2A1 is enriched at the centrosome, consistent with previous findings (25) (**Fig 2B**). We used 3D STED to examine endogenous CSNK2A1 localization relative to the centriolar core component Centrin-2, and the ciliary membrane protein ARL13B. This analysis revealed that CSNK2A1 predominantly localizes to the mother centriole at the vicinity of the axoneme (**Fig 2C**). CSNK2A1 signal shows a distinctive localization pattern, wherein it surrounds the mother centriole in a barrel-like fashion (**Fig 2C**). We distinguished two main loci of the CSNK2A1 signal. A symmetrical arc-like structure is found at the proximal end of the mother centriole. Fainter parallel linkers to centriolar side walls connect the proximal CSNK2A1 to a tighter distal locus (**Fig 2C, F**). In addition, we detect a weaker signal of the kinase at the daughter centriole (**Fig 2C**, **upper panel).**

To further characterize the localization of CSNK2A1 at the distal mother centriole, we analyzed its positioning to the distal appendage protein CEP164 together with γ-Tubulin and acetylated α-Tubulin using the structured illumination microscopy (SIM) (**Fig 2D**). In a side view, we observed an identical pattern to that obtained from co-labeling with Centrin-2. In contrast, in a top-down view, we noted a ring of CSNK2A1 partially overlaps with that of CEP164 (**Fig 2D, F**). Top view acquisition co-stained with TTBK2 confirmed partial co-localization between CSNK2A1 and TTBK2 rings at the mother centriole (**Fig 2E**) and indicates that CSNK2A1 is a centriolar distal appendage-associated protein (**Fig 2D, E, F**). Altogether, these results imply that CSNK2A1 and TTBK2 distal rings exhibit relatively similar positioning at the distal appendages, with evidence of a more complex organization for CSNK2A1 (**Fig 2F**).

We identified CSNK2A1 from a CRISPR screen for genes that modify the function of TTBK2, specifically as a gene that, when knocked down, increases the ability of hypomorphic *Ttbk2*^*null/gt*^ cells to activate a HH signaling response (**Fig 1B**). We previously demonstrated that *Ttbk2*^*null/gt*^ embryos have patterning defects due to reduced HH signaling, and a reduced frequency of cilia formation. Cells derived from these embryos similarly form cilia at a reduced frequency and these cilia have defects in IFT protein localization as well as structural abnormalities (9). Therefore, we tested whether the loss of CSNK2A1 function is sufficient to rescue or partially rescue cilia defects seen in *Ttbk2*^*null/gt*^ cells. To this end, we tested 5 different sgRNAs against *Csnk2a1*, from which we chose the most efficient sgRNA and selected individual clones to validate for CSNK2A1 knockdown by Western blot. We identified a clone for which *Csnk2a1* expression was entirely eliminated in *Ttbk2*^*null/gt*^ cells (**Fig S3A**).

Simultaneously, we isolated a control clonal cell line, generated by a non-targeting gRNA. Consistent with the identification of *Csnk2a1* in our CRISPR screen, sgRNA-mediated knockout of *Csnk2a1* in *Ttbk2*^*null/gt*^ cells partially rescued defects in cilia frequency as well as cilia length (**Fig 2G**).

### Loss of CSNK2A1 disrupts ciliary trafficking

To further examine the requirements for CSNK2A1 in cilia assembly and function, we generated clonal cell lines from WT MEFs that were CRISPR-engineered to deplete CSNK2A1 **(Fig S3B)**. Following 48 h of serum starvation to induce ciliogenesis, the frequency of cilia formation was not affected in the absence of CSNK2A1. However, *Csnk2a1*-depleted cells formed longer cilia on average with a wide range of measurements suggesting possible ciliary instability (**Fig 3A**).

**Figure 3:**
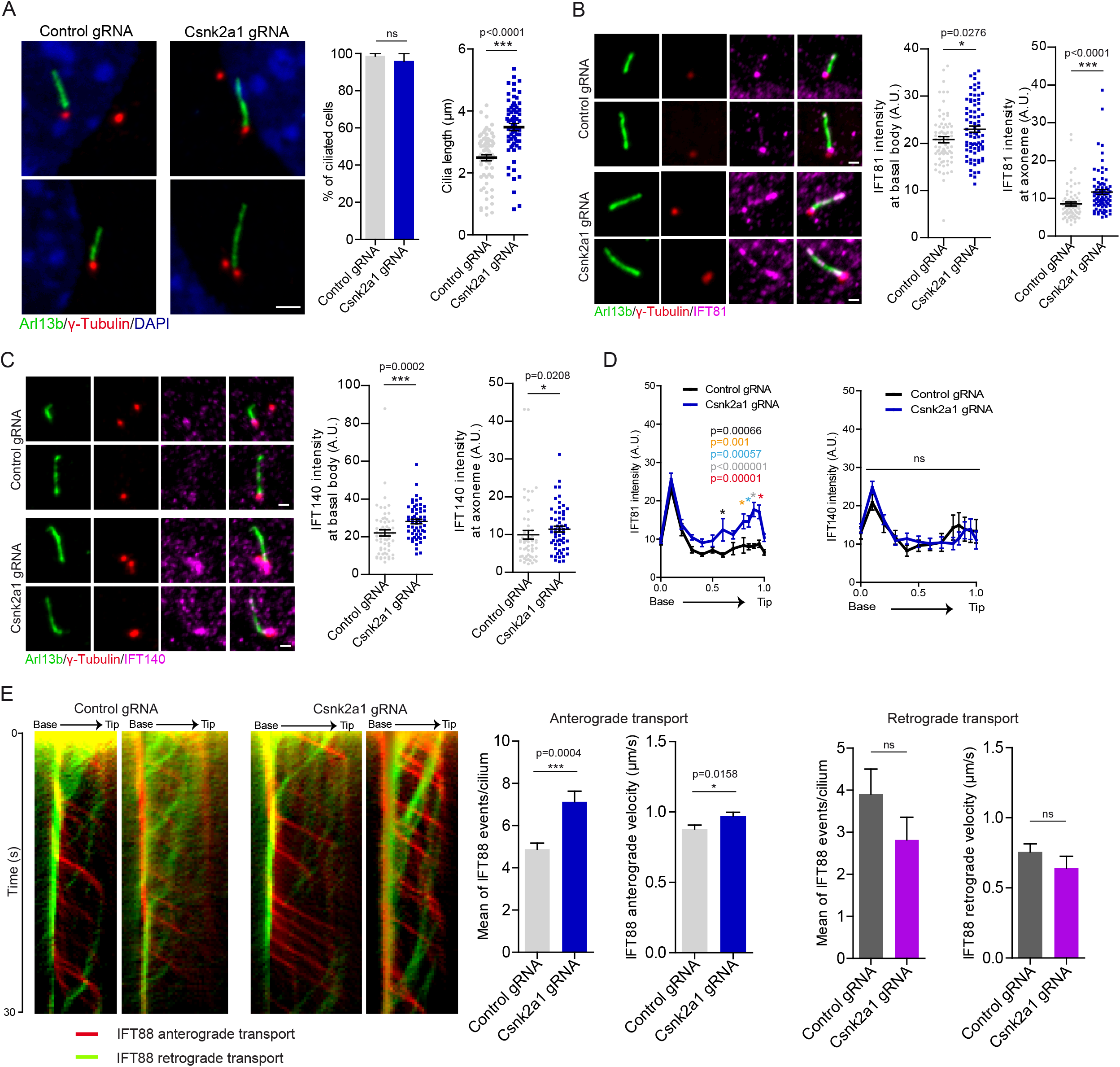
CSNK2A1 regulates cilia length and IFT trafficking. A. *Csnk2a1* knockout clonal cell line forms abnormally longer cilia. Representative immunofluorescence images of cilia, stained with antibodies to ARL13B (green) and γ-Tubulin (red), from control and *Csnk2a1* KO-1 clonal cells as indicated. Graphs show the mean percentage of ciliated cells and the average cilia length ± SEM. Statistical comparison was performed using the nonparametric Mann-Whitney test. P-values are displayed on the graph (ns: not significant). The total number of cells analyzed for both graphs is 69 cells per condition. Scale bar: 2 μm B. IFT81 accumulates at the basal body and cilia in *Csnk2a1* KO-1 cells. Representative immunofluorescence images of serum-starved cells are shown, labeled with the ciliary marker ARL13B (green), the centrosomal protein γ-tubulin (red), and IFT81 (magenta). Graphs show the mean intensity of IFT81 ± SEM at the basal body (left graph) and the axoneme (right graph) in control and *Csnk2a1* KO-1 cells. Statistical comparison was performed using the nonparametric Mann-Whitney test. P-values are shown on the graph. The cell number details presented in the graphs are as follows: Basal body: control gRNA n=72, *Csnk2a1* gRNA n=79 cells; Axoneme: control gRNA n=71, *Csnk2a1* gRNA n=79 cells. Scale bars: 1 μm. C. IFT140 accumulates in *Csnk2a1*-mutant cilia. Representative immunofluorescence images of serum-starved cells are displayed, stained with the ciliary marker ARL13B (green), the centrosomal protein γ-tubulin (red), and IFT140 (magenta). Graphs show the mean intensity of IFT140 ± SEM at the basal body (left graph) and the axoneme (right graph) in control and *Csnk2a1* KO-1 cells. Statistical comparison was performed using the nonparametric Mann-Whitney test. P-values are shown on the graph. Basal body: control gRNA n=58, *Csnk2a1* gRNA n=57 cells; Axoneme: control gRNA n=56, *Csnk2a1* gRNA n=57 cells. Scale bars: 1 μm. D. IFT81, but not IFT140, exhibits a ciliary distal accumulation in *Csnk2a1* KO-1 cells. The mean intensity of IFT81 (left graph) and IFT140 (right graph) ± SEM is profiled along the cilium. All cilia lengths were normalized to 1 and displayed on the X-axis. Statistical comparison was executed by multiple t-tests. P-values are shown on the graph (ns: not significant). Left graph: control gRNA n=28, *Csnk2a1* gRNA n=29 cells; Right graph: control gRNA n=30, *Csnk2a1* gRNA n=35 cells. E. The anterograde trafficking of IFT88-GFP, but not the retrograde, is increased in *Csnk2a1* KO-1 cells. Representative kymographs of IFT88-GFP for anterograde (red) and retrograde (green) trafficking are shown from the base to the ciliary tip for a duration of 30s (250 ms interval between two consecutive frames). Graphs exhibit the mean number of IFT88-GFP events per cilium and their velocities. A range of 2 to 5 values was extracted from each cilium. Error bars denote the SEM. Statistical comparison was performed using the nonparametric Mann-Whitney test. P-values are shown on the graph (ns: not significant). Total number of cilia quantified is as follows: Anterograde transport: Control gRNA n=18, *Csnk2a1* gRNA n=15 cilia; Retrograde transport: Control gRNA n=15, *Csnk2a1* gRNA n=13 cilia.

Cilia formation is a multistep process where TTBK2 plays an essential role (8). TTBK2 functions upstream of IFT proteins and triggers the final steps required for the conversion of the mother centriole into a functional basal body. As CSNK2A1 partially opposes TTBK2 function, we next evaluated IFT recruitment and localization in the cilia of *Csnk2a1*-knockdown cells. In contrast with *Ttbk2* mutant cells, *Csnk2a1*-knockdown cells have aberrant ciliary localization of components of the IFT-B complex that are critical for cilium assembly (26–28). Specifically, IFT81 and IFT88 abnormally accumulate in *Csnk2a1* mutant cilia, especially within the more distal axoneme (**Fig 3B, D**; **Fig S3C, D**). We also assessed the localization and intensity of IFT140, a component of the IFT-A complex. The IFT-A complex is implicated in retrograde transport as well as in the trafficking of specialized ciliary membrane proteins and HH pathway components (29, 30). In WT cells, IFT140 accumulates primarily at the ciliary base. In mutant cilia, we don’t observe substantial changes in its localization pattern in contrast with the IFT-B components, however, there is a modest increase in IFT140 intensity at the ciliary base and the axoneme in *Csnk2a1* knockdown cells (**Fig 3C, D**).

The changes we observed in IFT localization in *Csnk2a1*-depleted cilia, while modest, were highly reproducible. We postulated this may point to defects in intra-ciliary transport. To examine the bidirectional IFT transport, we monitored IFT88-GFP within the cilium by live imaging and measured anterograde and retrograde dynamics (**Fig 3E**). Mutant cilia had a greater number of IFT88 tracks moving towards the ciliary tip, with increased velocity speed compared to the control cilia. Retrograde measurements did not show a significant change compared to control cilia (**Fig 3E**). Altogether, these results suggest that CSNK2A1 regulates the appropriate levels of IFT recruited to the cilium.

### Trafficking of HH pathway components and signaling are disrupted in *Csnk2a1*-depleted cells

Given the disruptions in ciliary trafficking that we noted, we next assessed whether the trafficking of HH pathway components including SMO, GLI2, and KIF7 that localize to primary cilia is affected. Upon stimulation of HH signaling either with SHH ligand or with SMO agonist (SAG), the transmembrane protein SMO becomes enriched throughout the ciliary membrane (31). GLI2, a transcriptional activator of the pathway becomes enriched at the distal tip of cilia (32), as does the atypical kinesin KIF7, which regulates the structure of the ciliary tip compartment as well as participating in HH signaling (33, 34).

For each of these proteins, we observed that their accumulation in the cilium upon treatment with SAG was increased in *Csnk2a1*-depleted cells (**Fig 4A-C**). This increased accumulation or trafficking of HH pathway components into the cilium is often associated with changes in the transcriptional output of the pathway. To test whether this is the case for the *Csnk2a1*-depleted cells, we assessed the expression of two direct transcriptional targets of the HH pathway using qPCR: *Ptch* and *Gli1*. Consistent with the increased accumulation of HH pathway components within the *Csnk2a1* mutant cilia, we observed increased expression of both *Ptch* and *Gli1* upon SAG treatment in the mutant cells relative to WT (**Fig 4D**). In addition, we also noted that levels of *Ptch1* are higher in the *Csnk2a1*-mutant cells relative to WT even in the absence of SAG, suggesting that the basal levels of HH might be higher in the mutants (**Fig 4D**). Altogether, these results suggest that CSNK2A1 is a negative regulator of SHH pathway.

**Figure 4:**
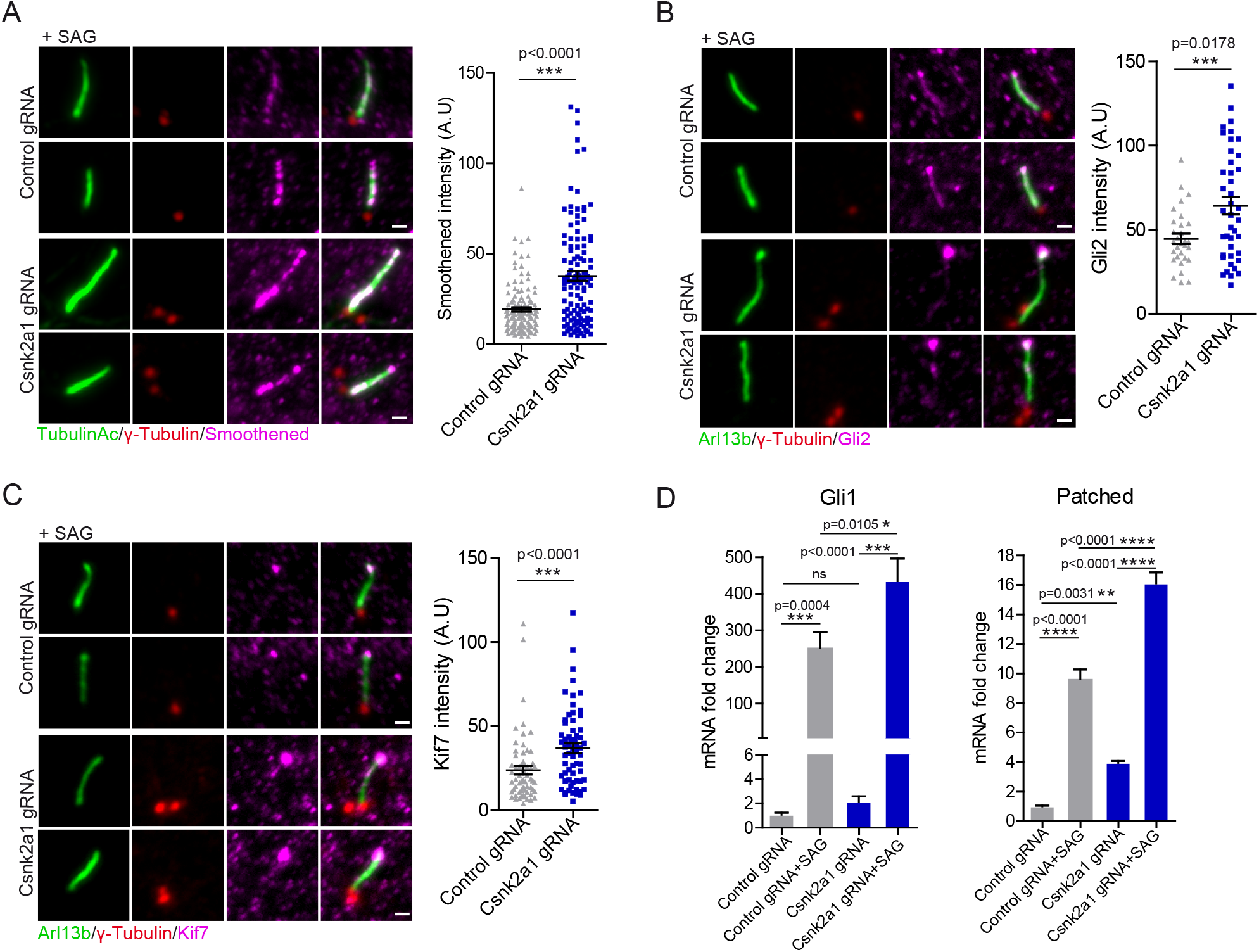
CSNK2A1-mutant cilia exhibit increased SHH response. A. SMO accumulates in *Csnk2a1-*mutant cilia. Cells were serum-starved in the presence of SAG. Representative immunofluorescence images of cilia, stained with antibodies to Acetylated Tubulin (green), γ-Tubulin (red) and SMO (magenta), from control and *Csnk2a1* KO-1 clonal cells. The graph shows the mean intensity of Smoothened ± SEM along the cilium in control and *Csnk2a1* KO-1 cells. Statistical comparison was performed using the nonparametric Mann-Whitney test. P-value is displayed on the graph. The data points were extracted from the following cell number: control gRNA n=123; *Csnk2a1* gRNA n=120 cilia. Scale bars: 1 μm. B. *Csnk2a1*-mutant cilia exhibit higher levels of GLI2 at the tip. Cells were serum-starved in the presence of SAG. Representative immunofluorescence images of cilia, labeled with antibodies to ARL13B (green), γ-Tubulin (red), and GLI2 (magenta), from control and *Csnk2a1* KO-1 clonal cells. Graph shows the mean intensity of GLI2 ± SEM at the ciliary tip in control and *Csnk2a1* KO-1 cells. Statistical comparison was performed using the nonparametric Mann-Whitney test. P-value is displayed on the graph. The data points were collected from control gRNA n=30 cilia, *Csnk2a1* gRNA n=41 cilia. Scale bars: 1 μm. C. *Csnk2a1*-mutant cilia show increased levels of KIF7 at the ciliary tip. Cells were serum-starved in the presence of SAG. Representative immunofluorescence images of cilia, labeled with antibodies to ARL13B (green), γ-Tubulin (red), and KIF7 (magenta), from control and *Csnk2a1* KO-1 clonal cells. Graph shows the mean intensity of KIF7 ± SEM at the tip in control and *Csnk2a1* KO-1 cells. Statistical comparison was performed using the nonparametric Mann-Whitney test. P-value is shown on the graph. Examined data points were collected from control gRNA n=63 cilia, *Csnk2a1* gRNA n=64 cilia. Scale bars: 1 μm. D. Real-time quantitative PCR shows a significant increase in mRNA fold change ± SEM of Gli1 and Patched transcripts in *Csnk2a1* KO-2 cells compared to control cells. Graphs were produced from 3 replicates per condition from two independent experiments. Statistical comparison was performed by one-way analysis ANOVA with Tukey’s multiple comparisons test. P-values are shown on the graph (ns: not significant).

### CSNK2A1 is required for cilium stability

Because the cilia of *Csnk2a1*-depleted cells are elongated and exhibit a variety of trafficking defects, we further examined the stability aspect of these cilia. Immunofluorescent staining of cilia revealed that *Csnk2a1*-depleted cells display a higher percentage of cilia with a tip gap that is positive for ARL13B but negative for acetylated α-Tubulin (**Fig 5A**). This gap, in addition to the accumulation of GLI2 and KIF7 in the tips of *Csnk2a1*-depleted cilia relative to WT, points to potential abnormalities in the ciliary tip compartment. To test whether defects at the ciliary tip were associated with decreased ciliary stability, we incubated the cells with nocodazole (Noco) to induce microtubule depolymerization, and to challenge the axonemal integrity of the cilium. We then calculated the ciliary length difference between the Noco-treated and the Noco-untreated conditions and noted that mutant cilia are less stable than control cilia (**Fig 5B**). Taken together, these data indicate that *Csnk2a1*-mutant cilia have an abnormal ciliary tip compartment that exhibits intrinsic structural instability.

**Figure 5:**
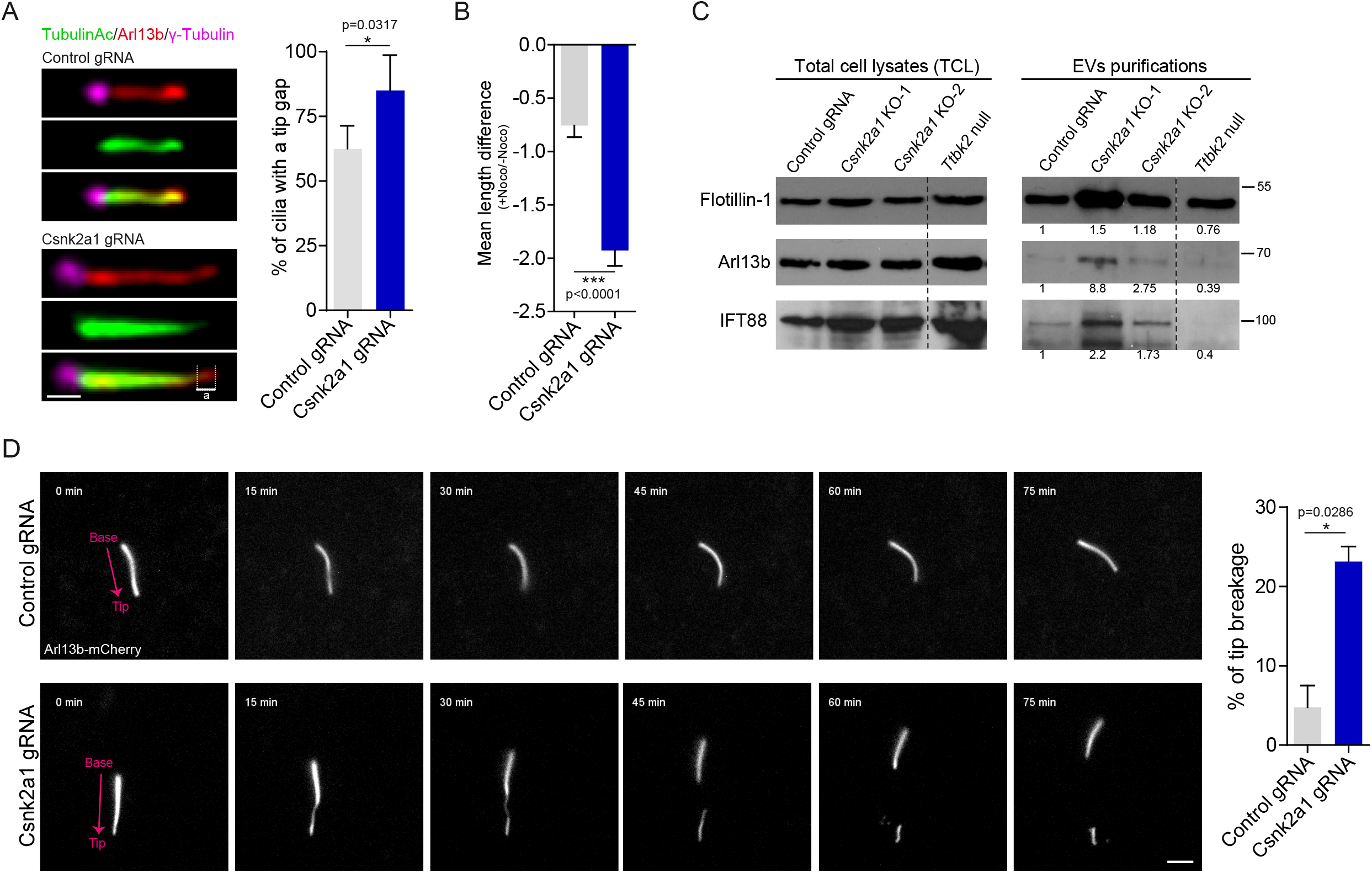
CSNK2A1 is essential for tip stability. A. The coverage of the ciliary tip by acetylated tubulin is reduced in *Csnk2a1* mutant cilia. The tip gap is a tip that is positive for ARL13B and negative for acetylated alpha-tubulin (illustrated by ‘’a’’). Scale bars: 1 μm. Graph shows the mean percentage of cilia that display tip gaps ± SEM (47 control cilia and 49 mutant cilia). B. Under Nocodazole (Noco) treatment, mutant cilia are less stable than control cilia. Cells were serum-starved for 48h and then treated with 10 μM Nocodazole or DMSO as a control for 35 minutes. Cilia length was collected from ARL13B immunofluorescent staining. Data is displayed as a mean difference of cilia length [+Noco]-[−Noco] ± SEM. Measurements were produced from 74 control cilia and 70 *Csnk2a1* mutant cilia. C. *Csnk2a1*-depleted cells shed more ciliary EVs. Western blots from total cell lysates (TCL, left blots) and EVs purifications (right blots) are probed with antibodies to FLOTILLIN-1, ARL13B, and IFT88. Two clonal *Csnk2a1*-depleted cell lines were used in comparison to control cells. *Ttbk2*^*null*^ cell line does not form cilia and serves as a negative control to evaluate the fraction of ciliary EVs. For each probe, the fold change value was calculated as follows: Ratio of [EVs_Area under the curve]: [TCL_Area under the curve]. The fold change was normalized to the control value. Dashed line marks where the western blot images were cut to remove experimental conditions that were not relevant to the current study. D. *Csnk2a1*-depleted cells are prone to tip breakage. Non-clonal control and *Csnk2a1*-depleted cells (50-60% of CSNK2A1 depletion by immunofluorescence) were generated in transgenic ARL13B-mCherry MEFs. Time-lapse imaging of cilia was carried out for 75 min. Breakage events were quantified and normalized to the total number of imaged cilia. The graph represents the mean percentage of tip breakage ± SEM for control (n=51 cilia) and *Csnk2a1*-depleted cells (n=63 cilia). Scale bars: 3 μm.

The ciliary tip is also a site of ectocytosis, a process by which vesicles are shed from the cilium. This process is linked to signaling from primary cilia and proper regulation of ectocytosis is important for controlling the ciliary length and for maintaining the stability of cilia (35–39). To test whether ciliary scission and ectocytosis could be affected in *Csnk2a1*-depleted cells leading to ciliary instability, we purified extracellular vesicles (EVs), that included cilia shedding through scission from supernatants collected from the culture media of control and *Csnk2a1*-mutant cells. The EVs fraction was collected through sequential ultracentrifugation, and subjected to lysis and western blotting for a marker of extracellular vesicles (FLOTILLIN) as well as ciliary proteins IFT88 and ARL13B. In total cell lysates, these proteins were found to be expressed at roughly equal levels between control cells and two different CRISPR-knockout clonal lines of *Csnk2a1* (**Fig 5C**). However, we noted that among the supernatants, we observed higher levels of FLOTILLIN as well as IFT88 and ARL13B in the *Csnk2a1* knockdown cells relative to controls. Additionally, we found that *Ttbk2* null mutant cells, which do not form primary cilia (8) have FLOTILLIN levels in the supernatant at levels comparable to WT cells, however, we observed a dramatic reduction of ciliary associated proteins IFT88 and ARL13B recovered in this fraction. Taken together, these data suggest that *Csnk2a1*-mutant cilia exhibit an increased amount of ciliary scission and/or EV formation.

To independently assess and extend these findings and examine how the loss of CSNK2A1 might affect cilium stability through ciliary scission, we depleted CSNK2A1 in a transgenic cell line expressing ARL13B-mCherry in order to perform live-imaging of cilia. Cells were then imaged every 15 minutes for 75 minutes. Interestingly, *Csnk2a1* mutant cilia frequently showed a thinning towards the distal part of the axoneme followed by a breakage of the ciliary tip. In comparison to control cells, mutant cilia showed an average increase of approximately 5-fold in the frequency of tip breakage/excision (**Fig 5D**, **V1-2 videos**).

Previously, actin and actin-binding proteins have been linked to ciliary scission (36, 37), similar to what we observed in our mutant cells. We, therefore, examined if our ciliary breaks were driven by changes in the actin cytoskeleton at the primary cilia. Live imaging of cells co-transfected with SSTR3-GFP and Lifeact-mCherry (F-actin) showed that F-actin was transiently recruited to the breakage site in *Csnk2a1* mutant cilia, prior to the excision event (**Fig S4A**, **V3-4 videos**). Similarly, induction of cilia disassembly by serum re-addition led to a distal enrichment of F-actin in *Csnk2a1*-depleted cilia (**Fig S4B**). Altogether, these results indicate that CSNK2A1 is required for the integrity of the ciliary tip and that the protein depletion causes spontaneous ciliary tip excision, likely driven by the actin cytoskeleton.

Dominant mutations in *Csnk2a1* were recently associated with a human syndrome known as Okur-Chung neurodevelopmental syndrome (OCNDS) (20, 40–44). This disorder is characterized by dysmorphic facial features as well as neurological phenotypes, bearing some similarities to human ciliopathies. To test whether the OCNDS-associated mutations are associated with defects in primary cilia, we stably expressed the wild-type version of CSNK2A1 as well as three different OCNDS-associated variants R80H, D156H, and R191Q (**Fig 6A**) (41, 43, 45) in WT MEFs. We previously showed that the GFP tagged-CSNK2A1^WT^ localized to the centrosome, similar to the endogenous protein **(Fig 2B)**. Interestingly, most *Csnk2a1* variants were still recruited to the centrosome, except for CSNK2A1^R80H^, which exhibited impaired recruitment to the centrosome **(Fig 6B)**. We next assessed the ciliation rate and found no significant difference between these cell lines **(Fig 6C)**. Because the primary phenotypes of *Csnk2a1-* depleted cilia are related to cilia stability, we examined cilia morphology across these cell lines expressing human disease-associated mutations. We found that each of the OCNDS-associated mutations significantly increased the frequency of cilia with morphological abnormalities, consistently between the three CSNK2A1 variants, and in comparison to the WT CSNK2A1 (**Fig 6D**). About 70% of these abnormalities include a bifurcated ciliary tip and/or membranous branch-like protrusions, located on the distal axoneme. An additional, 30% of these abnormal cilia showed a collapsed-like morphology, where ARL13B-positive cilia exhibited non-elongated bulgy and hollow curled structures (**Fig 6D)**.

**Figure 6:**
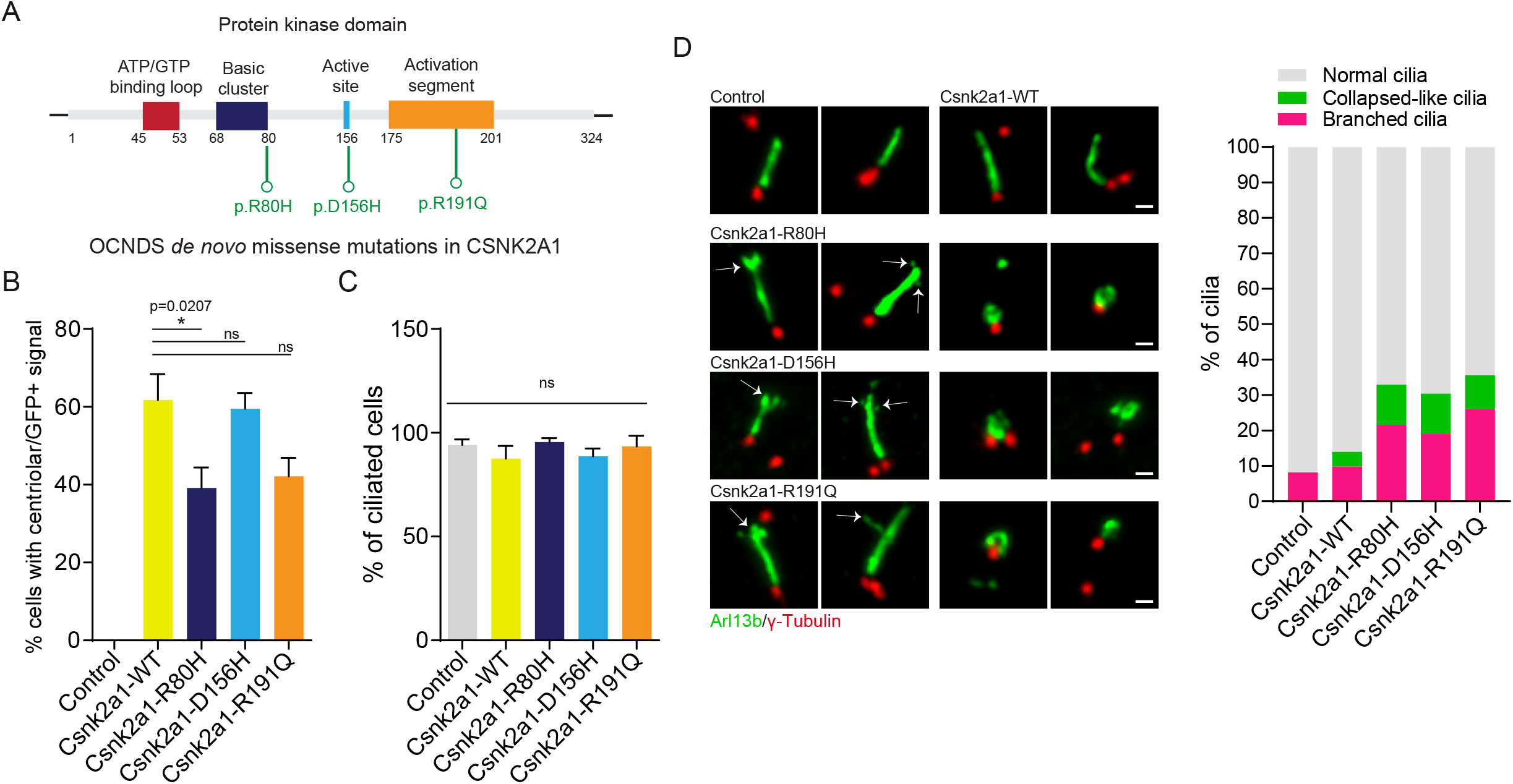
OCNDS-associated *Csnk2a1* mutations cause cilia structural defects. A. The kinase domain illustration of CSNK2A1 and the OCNDS-associated mutations. The catalytic subunit is composed of an ATP/GTP binding loop, adjacent to the basic cluster. The active site is located at the amino acid 156, close to the activation segment. The location of three OCND *de novo* missense mutations p.R80H, p.D156H, and p.R191Q are displayed in green. B. Graphs show the mean percentage of cells with centrosomal GFP-positive signal in control and CSNK2A1-expressing cell lines. Statistical comparison was performed by one-way analysis ANOVA with Tukey’s multiple comparisons test. P-values are shown on the graph (ns: not significant). Each graph displays results analyzed from control cells n=46; CSNK2A1-WT n=83; CSNK2A1-R80H n=64; CSNK2A1-D156H n=96; CSNK2A1-R191Q n=78 cells. C. Graphs show the mean percentage of cilia in control and CSNK2A1-expressing cell lines. Statistical comparison was performed by one-way analysis ANOVA with Tukey’s multiple comparisons test (ns: not significant). Each graph displays quantified from control cells n=46; CSNK2A1-WT n=83; CSNK2A1-R80H n=64; CSNK2A1-D156H n=96; CSNK2A1-R191Q n=78 cells. D. WT MEFs stably expressing CSNK2A1-R80H-GFP, CSNK2A1-D156H-GFP, and CSNK2A1-R191Q-GFP exhibit a higher percentage of abnormal cilia, which are defined by two structural phenotypes: (1) a distal membrane branching at the ciliary tip and (2) a collapsed-like ARL13B structure. Scale bars: 1 μm. The stacked graph shows the percentage of cilia in three categories: branched, collapsed-like and normal cilia. Control cells, n=41; CSNK2A1-WT, n= 127; CSNK2A1-R80H, n=60; CSNK2A1-D156H, n=107 and CSNK2A1-R191Q, n=85 cells. The percentage of each defect is detailed as follows: for the branching phenotype: Control: 8.09%, CSNK2A1-WT: 9.8%, CSNK2A1-R80H: 21,64%, CSNK2A1-D156H: 19% and CSNK2A1-R191Q: 26.03%; for the ciliary collapsed-like defect: Control: 0%, CSNK2A1-WT: 4.19%, CSNK2A1-R80H: 11.28%, CSNK2A1-D156H: 11.4% and CSNK2A1-R191Q: 9.52%. Statistical comparison was performed by one-way analysis ANOVA with Tukey’s multiple comparisons test on the mean percentage of abnormal cilia (Branched + collapsed-like phenotypes). P-values are detailed as follows: CSNK2A1-WT vs CSNK2A1-R80H, p-value=0.0102; CSNK2A1-WT vs CSNK2A1-D156H, p-value=0.0334; CSNK2A1-WT vs CSNK2A1-R191Q, p-value=0.0026.

These results indicate that the OCNDS-associated mutations may alter the morphology and structure of primary cilia. OCNDS exhibits overlapping clinical features with ciliopathies (20, 43, 46, 47). Our work, therefore, establishes a new link between *Csnk2a1* and cilia regulation and raises the possibility that OCNDS might be linked, at least partially, to disrupted cilia structure or function.

## DISCUSSION

Our present work identifies the CK2 catalytic subunit, CSNK2A1 as a negative regulator of TTBK2 and the SHH pathway. CSNK2A1 acts in opposition to TTBK2 function mediating cilia structure and trafficking. Our results reveal that CSNK2A1 and TTBK2 physically and functionally interact, likely at the distal appendages. In the absence of CSNK2A1, mutant cilia are longer and exhibit instability at the ciliary tip. These defects are also accompanied by increased anterograde trafficking and accumulation of IFT and SHH-related proteins within the cilium (**Fig 7**). In addition, the OCNDS-associated mutations cause ciliary abnormalities that may highlight the requirement for CSNK2A1 function in ciliary homeostasis, potentially disrupted in the human disease.

**Figure 7:**
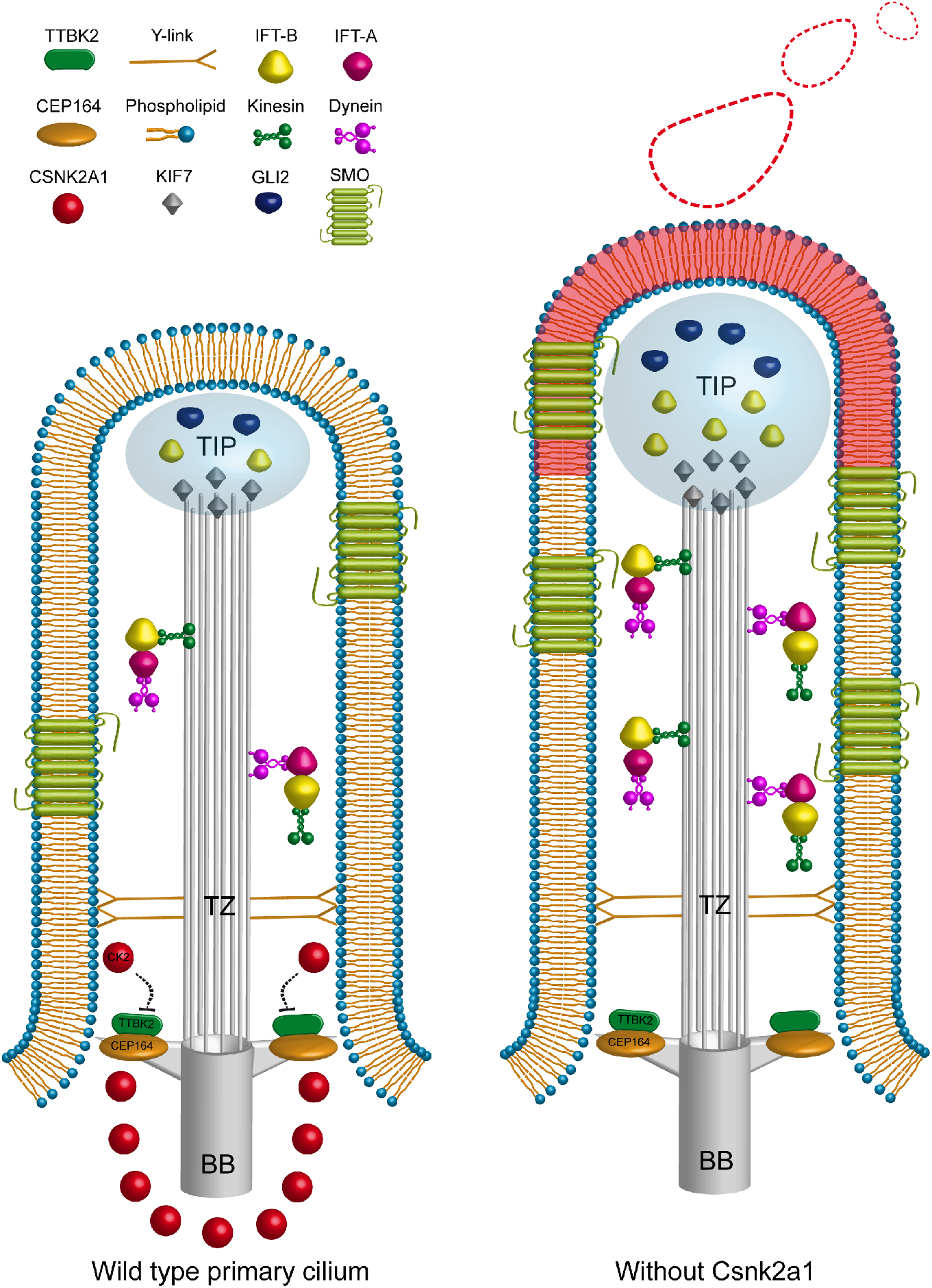
Working model of CSNK2A1 role in cilia homeostasis and function. CSNK2A1 acts as a negative regulator of TTBK2 and SHH signaling. It is differentially enriched at the mother centriole, forming a ring-like structure at the distal appendages. The kinase depletion leads to the aberrant trafficking of several critical regulators of cilia assembly (IFT proteins) and the SHH pathway that includes SMO, GLI2, and KIF7. CSNK2A1 mediates the proper influx of ciliary protein, implying a newly prominent role as a failsafe protein for SHH dysregulation.

The CK2 holoenzyme is comprised in most cases of two catalytic subunits (CSNK2A1 and/or CSNK2A2) and two regulatory CSNK2B subunits (14, 48, 49). CK2 plays a key role in numerous cellular processes, including signal transduction, cell cycle progression, and apoptosis (50–54). It is yet unclear whether the holoenzyme CK2 localizes to the centrosome and/or is implicated in ciliary function. Several reports showed that CK2 and the catalytic subunit CSNK2A1 may localize differentially in some contexts. For example, CK2 was found associated with the Golgi complex and the endoplasmic reticulum (55), whereas CSNK2A1 (McKendrick et al., 1999; This study) and CSNK2A2, but not CSNK2B, were detected at the centrosome (25). CSNK2A1 can also autonomously interact and/or phosphorylate an additional set of proteins, including PP2A, CALMODULIN, and MDM2 (57–60). Among the CK2 catalytic subunits, only CSNK2A1 came up as a hit in our TTBK2-related screens (This study). Taken together, it is conceivable that the individual catalytic subunit CSNK2A1 may function on its own in a localized manner at the centrosome and physically complexes with TTBK2, independently of the holoenzyme, to mediate cilia homeostasis.

Like TTBK2, we show that CSNK2A1 localizes to the distal appendages (DAPs), which are pinwheel-like structures emanating from the distal end of the mother centriole. DAPs function to control the membrane docking during ciliogenesis (61–64), and regulate the ciliary gate function (24). For example, the DAP protein FBF1 localizes to the distal end of the distal appendage matrix near to the membrane-docking site and is proposed to regulate the ciliary gating of transmembrane proteins, including Smoothened and SSTR3 (24). Similar to FBF1, loss of CSNK2A1 led to aberrant intra-ciliary levels of key proteins. Such trafficking defects may be caused by a dysfunctional ciliary gate that alters the amount of protein entry. Although we do not have a precise appreciation of the localization of CSNK2A1 relative to the ciliary membrane, the absence of the kinase may structurally affect the organization and function of the DAPs proximal to the ciliary membrane.

CSNK2A1 mutant cilia exhibit elevated levels of SMO, KIF7, GLI2, and IFT-B proteins along with a modest increase in SHH pathway activity. These defects partially overlap with those that characterize IFT-A complex mutants. For instance, *Ift122*^*null*^, *Ift139*^*null*^, and *Ift144*^*twt*^ (hypomorphic allele) cells accumulate IFT-B proteins, display aberrant retrograde transport, and show increased SHH activity (65–67). We speculate that the accumulation of ciliary proteins in these different mutants results from a disruption of the balance between anterograde and retrograde trafficking. For example, the retrograde trafficking is significantly slowed in *Ift139* mutants. In contrast, *Csnk2a1* mutant cilia exhibit increased anterograde trafficking both in terms of speed and number of events, which may cause molecular crowding at the ciliary tip. Thus, *Csnk2a1* compared to *IFT-A* mutant cilia exhibit striking similarities for aberrant ciliary contents, yet the cilia length phenotype appears distinct, which could be relative to differences seen in the dynamics of bidirectional transport.

Live imaging has revealed that cilia are dynamic organelles that shed ciliary vesicles in a phenomenon called cilia excision or decapitation. The release of ciliary vesicles is involved in the control of cell proliferation, cilia disassembly, and ciliary signaling (36–39). For example, when ciliary trafficking is perturbed, as in BBsome mutants, the rate of cilia excision increases as a mechanism to release the excess of GPCRs and prevent their accumulation in the axoneme (36). Our results indicate that increased anterograde transport in *Csnk2a1* mutant cilia correlates with high levels of IFT-B proteins at the ciliary distal end. This may eventually lead to a continuous lengthening of the cilium, triggering the ciliary excision release of the distal portion to the extracellular medium.

The ectopic ciliary tip excision might act as a failsafe mechanism, initiated to relieve and prevent a more severe accumulation of ciliary molecules at the ciliary tip. The mechanism by which CSNK2A1 mediates the stability of the ciliary tip is still unclear. It is possible that this kinase may regulate cilia stability and tip integrity by suppressing a key component implicated in cilia excision, such as the actin cytoskeleton (36, 37). Several studies reported that CK2 regulates the actin cytoskeleton dynamics (68–71). For example, CSNK2A1-dependent phosphorylation of CORTACTIN alters its ability to bind ARP2/3 and activate actin nucleation in the cytoplasm (71). In line with the literature, we found that the loss of CSNK2A1 triggered the abnormal formation of F-actin structures in cilia, prior to the scission. Hence, CSNK2A1 may function as a negative regulator of actin polymerization, restricting the formation of F-actin in cilia and preventing instability of the ciliary tip.

Recently, *de novo Csnk2a1* mutations were shown to cause Okur-Chung neurodevelopmental syndrome (OCNDS). Patients with OCNDS display a broad spectrum of clinical manifestations (20, 40–44) including cognitive impairment, hypotonia, skeletal defects, and ataxia, which overlap with phenotypical features of ciliopathies. The ciliopathies are a large group of pleiotropic disorders whose mutated genes are implicated in cilia function (20, 43, 46, 47), and are typically autosomal recessive. In contrast, OCNDS is autosomal dominant, with one mutated allele sufficient to cause the clinical features. Our result showed that stable expression of OCNDS-variants is sufficient to affect cilia morphology and structure, consistent with the dominant-negative nature of OCNDS mutations. Interestingly, the intriguing hollow structures that we noted in Fig 6D were similar to those described in *Bromi* mutant cilia (72), wherein the axonemes were curled, enveloped by dilated ciliary membranes, and exhibit mislocalized GLI2. This certainly raises the question of the effect of OCNDS mutations on cilia signaling and calls for future research to examine this key aspect.

Taken together, our present work identifies a novel kinase module CSNK2A1-TTBK2 that regulates cilia stability and signaling. The components of this module associate with the distal appendages and functionally opposes each other to mediate cilia function. CSNK2A1 may act as a negative modifier of TTBK2, regulating the trafficking and stability in cilia. Thus, our findings highlight the role of the distal appendage-associated kinase CSNK2A1 in ciliary trafficking, possibly through regulating the function of the ciliary gate. Our work also proposes a novel failsafe mechanism mediated by CSNK2A1 to prevent structural instability in cilia. We also find that OCNDS-associated mutations cause morphological defects in cilia that may relate to those we observe in *Csnk2a1* knockdown cells, suggesting a dominant-negative effect. Additional studies are needed to examine the molecular mechanisms by which *Csnk2a1* mutations trigger the developmental clinical features in OCNDS. Our work proposes that the pathologies of this disorder might be caused, at least in part, by dysfunctional ciliary signaling. Examining this possible link in detail and *in vivo* is therefore of high interest to us moving forward.

## Materials and Methods

### Cell culture and cell lines

Wildtype (WT), *Ttbk2*^*null/gt*^, *Ttbk2*^*null*^, and CRISPR-engineered mouse MEFs, and HEK293T cells were grown in high glucose DMEM (Gibco) with 10% FBS in a humidified 5% CO2 incubator at 37°C. Cilia formation was induced by serum starving the cells using media with 0.5% FBS. WT cell lines stably expressing TTBK2-FLAG-EGFP, CSNK2A1^WT^-FLAG-EGFP, CSNK2A1^R80H^-FLAG-EGFP, CSNK2A1^D156H^-FLAG-EGFP, and CSNK2A1^R191Q^ -FLAG-EGFP were maintained in the same growth media with 500 μg/ml of Geneticin.

### Constructs

To deliver sgRNAs into cells, we produced lentiviruses using pLentiCrisprV2 plasmid (52961, Addgene) (73). For each sgRNA, primers were designed, annealed, and inserted in the BsmBI site. For pLentiCrisprV2-non-targeting gRNA, the sgRNA sequence was 5’-ACCCATCGGGTGCGATATGG-3’. The CSNK2A1 sgRNA 5’-GGATTATGAATCACATGTGG-3’ was utilized to generate the pLentiCrisprV2-*Csnk2a1* sgRNA. The reference sequence of the mouse *Csnk2a1* cDNA was found in NCBI (NM_007788.3). We employed the Gateway destination vector pFLAP-Dest-EGFP-3xFLAG, where we introduced *Csnk2a1*^WT^ in the C-terminus of EGFP, separated by a linker of 67 amino acids. The site-directed mutagenesis method was utilized to generate pFLAP-DEST-CSNK2A1^R80H^-EGFP, CSNK2A1^D156H^-EGFP, and CSNK2A1^R191Q^ - EGFP. The following plasmids were also used in this study: pmCherry-pmCherry-Lifeact (Plasmid #54491, Addgene), pEGFPN3-SSTR3 (Plasmid #35623, Addgene), pCAGGS-mCherry-ARL13B (Goetz lab), and pCAGGS-EGFP-IFT88 (Goetz lab). BioID constructs for TTBK2 and GFP were generated using Gateway cloning into pDEST 5′ BirA*-FLAG-pcDNA5-FRT-TO (gift of Anne-Claude Gingras).

### Virus production and cell transduction

Lentiviruses were generated using a combination of three vectors. The sgRNA-Cas9 pLentiCrisprV2 construct, the packaging plasmid psPAX2, and the envelope plasmid pCMV-VSVg were transfected using polyethylenimine (7.5mM) in HEK293T cells for 20h. The transfection medium was removed and replaced with growth media. Virus-containing supernatant was collected 24h later and filtered through a 0.45 μm filter. For the production of retroviruses, we transfected pDEST-FLAP constructs with pCMV-VSVg in Phoenix cells, which stably express packaging proteins of Moloney Murine Leukemia Virus. Cells were transduced by the addition of viral supernatant in the growth medium in the presence of 8 μg/ml of polybrene. Following 24h incubation, the viral medium was replaced by the appropriate antibiotic-containing growth medium (5 μg/ml of Puromycin, 500 μg/ml of Geneticin). The selection of infected cells was carried out for 10 days.

### Generation of CSNK2A1 knockout cell lines

Wildtype and *Ttbk2*^null/gt^ MEFs were transduced with control or *Csnk2a1* sgRNA lentiviruses. At 24 hours post-infection, cells were selected with 5 μg/ml puromycin-containing media. Clonal cell lines were generated by limiting dilution. Clones were picked and grown in 24-well plates. For each clone, we prepared a total cell lysate for western blot analysis. We only retained *Csnk2a1*-depleted clonal cell lines that showed a complete knockout.

### BioID of TTBK2

#### Generation of stable cell lines for biotin proximity labeling

HEK293T cells stably expressing a single copy of N-terminally tagged TTBK2- or GFP-BirA* were generated using Flp-IN T-REx cells (ThermoFisher K6500-01), Positive clones were selected with hygromycin resistance, and maintained in media with tetracycline-free FBS.

For the proximity labeling experiments, cells were cultured in serum-free media for 48 hours to induce ciliogenesis, with tetracycline (1μg/ml) added to induce expression of the BirA* fusion constructs. Biotin (50μM) was then added to the cultures for an additional 24 hours to induce labeling. Cell were lysed in 50mM Tris pH 7.5, 100mM NaCl, 0.5% NP-40, and 10% glycerol, and briefly sonicated. Cleared lysates were incubated with washed Neutravidin beads, 4°C rotating, for 4 hours. Beads were then washed in 50mM ammonium bicarbonate (pH8.3) and submitted to the Duke University Proteomics and Metabolomics Core Facility for further processing and Mass spectrometry.

#### Sample Preparation

6 samples were submitted to the Duke Proteomics Core Facility (DPCF) (3 replicates of both TTBK2 and GFP). Samples were supplemented with 40μL 10% SDS in 50 mM TEAB, then reduced with 10 mM dithiolthreitol for 30min at 80C and alkylated with 20 mM iodoacetamide for 30min at room temperature. Next, they were supplemented with a final concentration of 1.2% phosphoric acid and 683 μL of S-Trap (Protifi) binding buffer (90% MeOH/100mM TEAB). Proteins were trapped on the S-Trap, digested using 20 ng/μl sequencing grade trypsin (Promega) for 1 hr at 47C, and eluted using 50 mM TEAB, followed by 0.2% FA, and lastly using 50% ACN/0.2% FA. All samples were then lyophilized to dryness and resuspended in 12 μL 1%TFA/2% acetonitrile containing 12.5 fmol/μL yeast alcohol dehydrogenase (ADH_YEAST). From each sample, 3 μL was removed to create a QC Pool sample which was run periodically throughout the acquisition period.

#### Quantitative Analysis, Methods

Quantitative LC/MS/MS was performed on 4 μL of each sample, using a nanoAcquity UPLC system (Waters Corp) coupled to a Thermo QExactive HF-X high-resolution accurate mass tandem mass spectrometer (Thermo) via a nanoelectrospray ionization source. Briefly, the sample was first trapped on a Symmetry C18 20 mm × 180 μm trapping column (5μl/min at 99.9/0.1 v/v water/acetonitrile), after which the analytical separation was performed using a1.8 μm Acquity HSS T3 C18 75 μm × 250 mm column (Waters Corp.) with a 90-min linear gradient of 5 to 30% acetonitrile with 0.1% formic acid at a flow rate of 400 nanoliters/minute (nL/min) with a column temperature of 55C. Data collection on the QExactive HF mass spectrometer was performed in a data-dependent acquisition (DDA) mode of acquisition with a r=120,000 (@ m/z 200) full MS scan from m/z 375 – 1600 with a target AGC value of 3e6 ions followed by 30 MS/MS scans at r=15,000 (@ m/z 200) at a target AGC value of 5e4 ions and 45 ms. A 20s dynamic exclusion was employed to increase depth of coverage. The total analysis cycle time for each sample injection was approximately 2 hours. Following 8 total UPLC-MS/MS analyses, data was imported into Proteome Discoverer 2.2 (Thermo Scientific Inc.), and analyses were aligned based on the accurate mass and retention time of detected ions using Minora Feature Detector algorithm in Proteome Discoverer. Relative peptide abundance was calculated based on area-under-the-curve (AUC) of the selected ion chromatograms of the aligned features across all runs. The MS/MS data was searched against the SwissProt H. sapiens database (downloaded in Nov 2017) with additional proteins, including yeast ADH1, bovine serum albumin, as well as an equal number of reversed-sequence “decoys”) false discovery rate determination. Mascot Distiller and Mascot Server (v 2.5, Matrix Sciences) were utilized to produce fragment ion spectra and to perform the database searches. Database search parameters included fixed modification on Cys (carbamidomethyl) and variable modifications on Meth (oxidation) and Asn and Gln (deamidation). Peptide Validator and Protein FDR Validator nodes in Proteome Discoverer were used to annotate the data at a maximum 1% protein false discovery rate.

### Kinome CRISPR-Cas9 suppressor screen of TTBK2

#### Lentiviral production

In this screen, we used a mouse pooled kinome CRISPR-Cas9 pooled library (#75316, Addgene) (23), which contains 2851 unique sgRNAs targeting 713 mouse kinase genes. It also includes 100 non-targeting sgRNAs, as an internal control. The lentiviral library was produced in HEK293T cells, grown in 15cm plates. We transfected the following plasmids sgRNA-Cas9 pLentiCrisprV2 (the library constructs), psPAX2, and pCMV-VSVg (3:2:1, respectively) for 20h using polyethylenimine (7.5mM). The transfection medium was removed and replaced by growth media. 48h later, virus-containing supernatants were collected and filtered using 0.45 μm filters. The viral titer was evaluated by transducing cells with a range of viral dilutions. Cells were then selected with 5 μg/ml puromycin for 48h. We employed the CellTiter-Glo Luminescent Cell Viability Assay (Promega, G7570) to determine the infectious units per ml (IFU/ml).

#### Kinome library in SHH-BlastR MEFs

This screen is based on the response of the Sonic hedgehog pathway. We introduced the transcriptional SHH-BlastR reporter (pGL-8xGli-Bsd-T2A-GFP-Hyg, (22)) in WT and *Ttbk2* hypomorphic (bby/GT) MEFs. The reporter was transfected in these cells that were selected and grown in media containing 200 μg/ml of hygromycin. The next step was to integrate the lentiviral library in SHH-BlastR MEFs. Cells were grown in 6-well plates and transduced at a multiplicity of infection of 0.2 in sufficient numbers such a manner that there was a 650:1 ratio of transduced cells per sgRNA. Cell infection was facilitated by centrifugation at 2200 rpm for 45 min at 25°C. The viral supernatant was incubated for 24h in the presence of 8 μg/ml polybrene. Cells were selected for 13 days in growth 10% FBS media with 200 μg/ml of hygromycin, and 5 μg/ml puromycin.

#### Kinome-Blasticidin reporter screening

SHH-BlastR MEFs were seeded in 15cm plates at a confluence of about 90%. For the screen, we maintained a 1700:1 ratio of cells to sgRNAs. Each experimental condition was achieved in duplicate for each cell line. Cells were serum-starved for 30h with or without 200nM SAG in 0.5% FBS media supplemented with 200 μg/ml of hygromycin, and 5 μg/ml puromycin. At this point, T0 cells were collected, and corresponding pellets were frozen at −20°C. The remaining plates were treated or not with 2.25 μg/ml blasticidin for 16h. Cells were then washed with 1x PBS, and growth media was added with or without 200nM of SAG. Media was changed every 2-3 days up until most cells in control plates (-SAG, +Blasticidin) stop cycling and die. WT and *Ttbk2* hypomorphic SHH-BlastR cell lines were collected at 4 days and 11 days, respectively, post-blasticidin treatment.

#### Sample preparation and Illumina sequencing

From the collected pellets, genomic DNA was extracted and purified using the DNeasy Blood and tissue kit (Qiagen). We amplified by PCR the integrated sgRNAs using P5/P7 primers, following the CRISPR library-related instructions (#75316, Addgene). All PCRs were pooled and sequenced by Illumina sequencing (Duke Center for Genomic and Computational Biology (GCB), Duke University).

#### Analysis of data from Illumina sequencing

Raw data from Illumina sequencing were converted to the number of reads for each individual sgRNA. A read was assigned to a specific target sequence if it matched the reference sequence with at most one mismatch (GCB, Duke University).

#### Analysis of sgRNA reads

All the counts per sgRNA were normalized with the total number of counts per condition using R studio. We then generated the ratio +blasticidin(+SAG): -Blasticidin(+SAG) for each sgRNA. Four sgRNAs represented each gene in the library. The average of the two median values for each gene was calculated to evaluate the gene depletion overall effect on the SHH response.

### Antibodies

Antibodies and their condition of use are detailed as follows: Anti-acetylated tubulin (Immunofluorescence (IF): 1:1000; Sigma, T6793); Anti-γ-Tubulin (IF: 1:500; Sigma, SAB4600239): Anti-Centrin2 (IF: 1:500; Millipore, 04-1624); Smoothened (IF: 1:500; Pocono Rabbit Farm and Laboratory, Inc.); Anti-ARL13B (IF:1/500, western blot (WB): 1/1000; NeuroMab, 11000053); Anti-CSNK2A1 (IF and western blot (WB): 1:500; Proteintech, 10992-1-AP); Anti-IFT88 (IF and WB: 1:500, Proteintech, 13967-1-AP); Anti-IFT81 (IF: 1:300, Proteintech, 11744-1-AP,); Anti-IFT140 (IF: 1:200, Proteintech, 17460-1-AP),; Anti-Flotillin-1 (WB: 1:1000, ABclonal, A6220); Anti-KIF7 (IF: 1/500; a gift from Dr. Mu He, (34)); Anti-GLI2 (IF: 1:500, a gift from J. T. Eggenschwiler); Anti-CEP164 (IF: 1:200, Sigma); Anti-FLAG M2 (IF: 1:75, WB: 1:4000, Sigma, F1804); Anti-GFP (IF: 1:500, WB: 1:1000, Proteintech, 66002-1-Ig); Anti-Vinculin (WB: 1/10000, Sigma, V4505).

### Lattice SIM

WT MEFs were plated on high precision type 1.5 coverslips and incubated in 0.5% serum media for 48h. Cells were fixed with cold methanol for 4.5 min. For a better signal, we incubated primary antibodies for 20h at 4°C. We used a combination of three secondary antibodies for each staining (IF: 1:500; Alexa Fluor 488, Alexa Fluor 568, Alexa Fluor 647; Molecular probes, Thermo Fisher Scientific). These were incubated for 2h at room temperature. Coverslips were then mounted on glass slides using ProLong Gold mounting media (Molecular probes, Thermo Fisher Scientific). SIM acquisitions were performed on a Zeiss Elyra 7 system equipped with lattice SIM technology. The signal was collected using a Zeiss Plan-Apochromat 63/1.4 Oil objective. The laser power and exposure times were kept at a minimum. In ZEN imaging software, the SIM mode, in combination with the tiling functionality, was used to process the acquired images.

### STED

Cells were prepared similarly to the Lattice SIM protocol. Instead, with two-colored STED, we labeled cells with Alexa Fluor 594 (Thermo Fisher Scientific) and ATTO 647N Antibodies (Sigma). STED was performed on a Leica DMi8 motorized inverted microscope coupled with a STED depletion laser (775 nm) and a pulsed white light laser (470nm-670nm). A Leica 100x/1.4 HCX PL APO oil objective was used to image ciliary and centrosomal labeling. 3D z-stacks were sequentially collected for each fluorophore using high sensitivity GaAsP Hybrid Detectors (HyDs) with gating capabilities for STED. Images were displayed and analyzed on the LAS X software. An additional STED deconvolution step was implemented in Huygens deconvolution software, linked to the LAS X software, to reach the optimal resolution.

### Ciliary vesicle purification

CRISPR-engineered cell lines and ttbk2^*null*^ MEFs were plated on 15cm dishes to reach a 90% confluence. Cells were serum-starved for 24h and subsequently washed 3 times with PBS to remove non-ciliary vesicles present in the media. Fresh 0.5% FBS media was added and incubated for 24h. Vesicles-containing supernatant was transferred to a 50ml conical tube. Cells were washed with 10ml PBS, then added to the same conical. We also prepared total cell lysates by lysing the cells directly in a 2X sample buffer. The conical tubes were centrifuged at 300 RCF for 10 min at 4°C. The supernatant was then transferred to a new 50ml conical and centrifuged for 20 min at 2000 RCF (4°C). The resulting supernatant was centrifuged for 40 min at 7500 RPM in an sw32 ti Rotor at 4°C. The supernatant was finally ultracentrifuged for 90 min at 25000 RPM in the same rotor at 4°C. The pellet was washed with 40 ml PBS and ultracentrifuged for 90min at 25000 RPM at 4°C. Most of the supernatant was then aspirated, and the remaining volumes were carefully equalized between samples. The sample buffer was finally added for western blotting analysis.

### Live imaging of cilia

Cells were seeded in 8-well Nunc Lab-Tek II chambered coverglass and serum-starved for 24h. At all times, cells were kept in a humidified chamber at 37°C. The live imaging was performed using a Zeiss Axio Observer-Z1 widefield microscope. The Zeiss Plan-Apochromat 63x / 1.4 Oil objective with the Axiocam 506 monochrome camera collected the fluorescent signal. The Zeiss definite focus was employed to maintain cilia on focus. Time-lapse series were generated using the Fiji software.

### IFT88 transport measurements

Cells were co-transfected with pCAGGS-mCherry-ARL13B and pCAGGS-EGFP-IFT88 and serum-starved for 24h. Cilia were live imaged every 250 ms for 30 sec to generate fast time-lapse series. These image sequences were converted into an 8-bit format. The unspecific background was removed with the correction of the unwanted cilia movement. IFT88 bidirectional transport along the cilium was displayed using kymographs that were generated using a Fiji macro toolset called KymographClear 2.0 (74).

### Immunofluorescence microscopy

Cells were plated on glass coverslips with an 80% confluence, followed by a serum starvation step of 48h to induce ciliogenesis. Fixation was carried out with 4% paraformaldehyde for 5 min and permeabilized with 100% cold methanol for 5 min. Fixed cells were blocked in blocking buffer (2% FBS, 0.1% Triton X-100 in PBS) for 1h and then incubated with primary antibodies for 3h at room temperature, followed by 2h incubation with secondary antibodies (1:500; Alexa Fluor 488, Alexa Fluor 568, Alexa Fluor 647; Molecular probes, Thermo Fisher Scientific). DNA was stained with DAPI for 1 min. Coverslips were mounted on glass slides using ProLong Gold mounting media (Molecular probes, Thermo Fisher Scientific). Cells were imaged with a Zeiss Axio Observer-Z1 widefield microscope equipped with a Zeiss Plan-Apochromat 63x / 1.4 Oil objective, an Axiocam 506 monochrome camera, and an Apotome setting. To accurately quantify the fluorescence intensities of ciliary markers, we acquired a Z-stack and created a maximum intensity projection for each image using Fiji software.

### Ciliation rate and fluorescence intensity quantification

Measurements of cilia rate and fluorescence intensity were performed from at least five randomly selected fields of cells for each condition. The fluorescence intensity of centrosomal and ciliary markers was quantified using the Fiji software. The polygon tool was chosen to select the positive pixels of the immunostained protein. We subsequently measured the mean fluorescence intensity, which integrates the total fluorescence intensity by the selected surface area.

### Western blot analysis

Cells were plated in a 6-well plate to reach a 90% confluence. Cells were directly lysed in 2x Laemmli sample buffer (50 mM Tris-HCl (pH 6.8), 10% glycerol, 2% SDS, 1% 2-mercaptoethanol, 0.1% bromophenol blue) on ice. Whole-cell extracts were incubated at 95°C for 5 min to denature proteins. Total cellular proteins were separated by SDS-PAGE and transferred onto a PVDF membrane. The membrane was saturated in 5% milk PBST (5% dry powdered milk, 0.1% Tween-20, 1X PBS) and incubated in primary antibodies followed by peroxidase-conjugated affinipure secondary antibodies. Membranes were developed with an ECL reagent kit (BioRad). Western blots were quantified using Fiji software (“Gels” command).

### Pulldowns

In the HEK293T cells, we transfected the pDEST40-TTBK2-V5 construct with pEGFPN3 or pFLAP-DEST-CSNK2A1^WT^-EGFP plasmids using polyethylenimine (7.5mM) for 16h. Transfection media was replaced with pre-warmed 10% serum media. We then serum-starved the cells for 72h in 0.5% serum media. Cells were collected on ice in 0.5% NP40 lysis buffer (50 mM tris-Hcl pH=8. 100 mM NaCl, 10% glycerol, EDTA-free protease inhibitor cocktail (Sigma), 10 μM of MG-132 (Sigma, M7449), 1mM EDTA, 10 mM N-Ethylmaleimide (Sigma, E3876) and 25 mM β-Glycerol phosphate). The cell lysate was incubated on ice for 10 min and then sonicated twice for 3 seconds with a 25% amplitude. We centrifuged the lysates for 20 min at 15000 rpm and recovered the supernatant. To pull down GFP-recombinant proteins, we incubated the GFP-Trap agarose beads (Chromotek, gta-10) with the total cell lysates for 3h at 4°C. These beads were previously blocked in 200 mg/ml BSA for 10 min at 4°C.

The beads were then collected by centrifugation (2500 rpm, 1 min) and washed five times with cold lysis buffer. After adding the sample buffer, samples were resolved by western blot. The transfer onto PVDF membrane was carried out for 5h at 4°C and at a current power of 80 Volt.

### RT-qPCR

Control and *Csnk2a1* knockout cells were plated in a 6-well plate. Cells were serum-starved for 48h and treated or not with 200 nM of SAG. Total RNA was purified using the RNeasy kit (Qiagen). From each condition, a total of 600 ng RNA was reverse transcripted to cDNA using the iScript Reverse Transcription Supermix for RT-qPCR (#1708840). Real-time quantitative PCRs were performed following the PowerUp SYBR Green (ThermoFisher) manufacturer instructions. We used the following primers: GLI1 forward 5’-TTATGGAGCAGCCAGAGAGA-3’, reverse 5’- GAGCCCGCTTCTTTGTTAAT-3’; PTCH1 forward 5’- TGACAAAGCCGACTACATGC- 3’, reverse 5’- AGCGTACTCGATGGGCTCT3’; ACTIN forward 5’- TGGCTCCTAGCACCATGA-3’, reverse 5’- CCACCGATCCACACAGAG-3’.

### Statistical analysis

Data are reported as arithmetic means ± SEM. Statistical analyses were performed using the nonparametric Mann-Whitney test with GraphPad Prism 8 software. For multiple comparisons, we executed a one-way analysis ANOVA with Tukey’s multiple comparisons test. To analyze the intensity profile data, we implemented multiple t-tests (one per row), found in the grouped analyses tab in GraphPad Prism 8 software. P ≤ 0.05 was used as the cutoff for statistical significance.

## Acknowledgments

We thank Drs. Kris Wood and Kevin Lin for their guidance with the CRISPR screen. We are grateful to Drs. David Breslow, Anne-Claude Gingras, Laurence Pelletier, and Saikat Mukhopadhyay for generously sharing reagents. We also thank Dr. Erik Soderblom and the Duke Proteomics and Metabolomics Core for their help with the TTBK2 BioID. Many thanks to Dr. Oliver Tress and Kathryn Schallhorn for acquiring the SIM data. This work was supported by the National Institutes of Health (R01 HD099784 to SCG), and the Regeneration Next Initiative at Duke University (RNI Postdoctoral Fellowship to AL).

## Supplementary figures

Figure S1:

A. TTBK2-BirA* expression rescues cilia defects in T*tbk2*^*null*^ MEFs. Representative immunofluorescence images were labeled with the ciliary marker ARL13B (green) and the centrosomal protein γ-tubulin (red). DNA was stained with DAPI.

B. Illustration of the experimental procedure of TTBK2 BioID. BirA*-GFP or BirA*-TTBK2 were introduced and stably expressed in Tet-ON Flp-In HEK293T cells. While BirA-GFP was homogenously expressed throughout the cell, BirA*-TTBK2 localized to the basal body in ciliated cells. These cell lines were serum-starved for 48h in the presence of tetracycline. Biotin was then added to cells and incubated for 24h. In cells expressing BirA2*-TTBK2, BirA* biotinylates known and unidentified interactors of TTBK2, most likely localized to the basal body. However, cells expressing BirA*-GFP will display more homogenous protein biotinylation throughout the cell. Biotinylated proteins were purified from total cell lysates and sequenced by mass spectrometry. Statistical analysis was conducted to identify potential interactors of TTBK2.

C. Purification of biotinylated proteins from inducible cell lines for GFP-BirA* or TTBK2WT-BirA2*. Cells were incubated with biotin and induced or not with Tetracycline. Pulldowns were performed with streptavidin beads in whole-cell extracts. Dashed line marks where the western blot image was cut to remove lanes, not relevant to this study.

Figure S2:

A. Illustration of the experimental procedure of the kinome CRISPR-Cas9 screen of TTBK2. The detailed description is found in the “materials and methods” section.

B. Same as figure 1B but for WT MEFs.

Figure S3:

A. Control or *Csnk2a1* gRNAs were used to generate clonal cell lines in *Ttbk2*^*null/gt*^ MEFs. *Csnk2a1* depletion was evaluated in total cell lysates from control and *Csnk2a1* KO cells, using western blot analysis. Vinculin was used as a loading control.

B. Western blots displaying CSNK2A1 depletion in two clonal cell lines generated in WT MEFs. Non-targeting gRNA was used to produce the control cell line. Blots are revealed with antibodies to CSNK2A1 and VINCULIN, shown as a loading control.

C. IFT88 accumulates at the basal body and cilia in *Csnk2a1* KO-1 cells. Representative immunofluorescence images of serum-starved cells are shown, labeled with the ciliary marker ARL13B (green), the centrosomal protein γ-tubulin (red), and IFT81 (magenta). Graphs show the mean intensity of IFT881 ± SEM at the basal body (left graph) and the axoneme (right graph) in control and *Csnk2a1* KO-1 cells. Statistical comparison was performed using the nonparametric Mann-Whitney test. P-values are shown on the graph. Basal body: control gRNA n=73 cells, *Csnk2a1* gRNA n=77 cells; Axoneme: control gRNA n=73 cells, *Csnk2a1* gRNA n=77 cells. Scale bars: 1 μm.

D. IFT88 exhibits a distal accumulation in *Csnk2a1* KO-1 mutant cilia. The mean intensity of IFT88 ± SEM is profiled along the cilium. All cilia lengths were normalized to 1 and displayed on the X-axis. Statistical comparison was executed by multiple t-tests. P-values are shown on the graph. Control gRNA n=20 cells and *Csnk2a1* gRNA n=20 cells.

Figure S4:

A. F-actin transiently localizes to mutant cilia prior to cilia excision. SSTR3-GFP and Lifeact-mCherry were overexpressed in control and *Csnk2a1* KO-1 cell lines. Time-lapse imaging was performed on ciliated cells for 40 min (1 acquisition every 10min). Scale bars: 2 μm.

B. F-actin is enriched in mutant cilia during cilia disassembly. SSTR3-GFP and Lifeact-mCherry were transfected in control and *Csnk2a1* KO-1 cells. Serum starvation was performed for 48h followed by serum re-addition to disassemble cilia. Cells were fixed and imaged with no immunostaining. Graphs show the intensity profiles of SSTR3-GFP (green) and Lifeact-mCherry (red), which display enrichment of Lifeact-mCherry in mutant cilia. Control cilia do not exhibit an apparent overlapping signal between the two markers. Scale bars: 1 μm.

V1 video: Transgenic ARL13B-mCherry control MEFs show normal cilia movement behavior.

V2 video: Transgenic ARL13B-mCherry *Csnk2a1*-depleted MEFs exhibit a tip breakage event. For V1 and V2 videos, cells were serum-starved to induce cilia formation. Frames of cilia were acquired every minute. The time-series are shown at a speed of 24 frames per second (fps).

V3 video: F-actin is not enriched in control cilia

V4 video: F-actin transiently localizes to cilia tip prior to breakage event in *Csnk2a1* KO-1 cells. For V3 and V4 videos, control and *Csnk2a1* KO-1 cells were transfected with Lifeact-mCherry and SSTR3-GFP and serum-starved to induce ciliogenesis. Acquisitions were taken every 10 min. Videos are shown at a speed of 0.5 fps.

Supp. Table 1: Results of TTBK2 BioID. Sheet 1 displays the raw data that was normalized to generate fold change and significance values. Sheet 2 shows the high confidence hits enriched in the TTBK2 condition in comparison to GFP.

Supp. Table 2: CRISPR-Cas9 data sets. Sheet 1 shows the raw counts for each gRNA. Each gene was targeted by 4 different gRNAs. Sheet 2 displays the mean of the medians for each gene from WT MEFs. Sheet 3 shows the mean of the medians for each gene from *Ttbk2* hypomorphic MEFs.

