## Supplementary figures and images for "A complex of distal appendage-associated kinases linked to human disease regulates ciliary trafficking and stability"

### Supplementary Figure S1

Figure S1

A

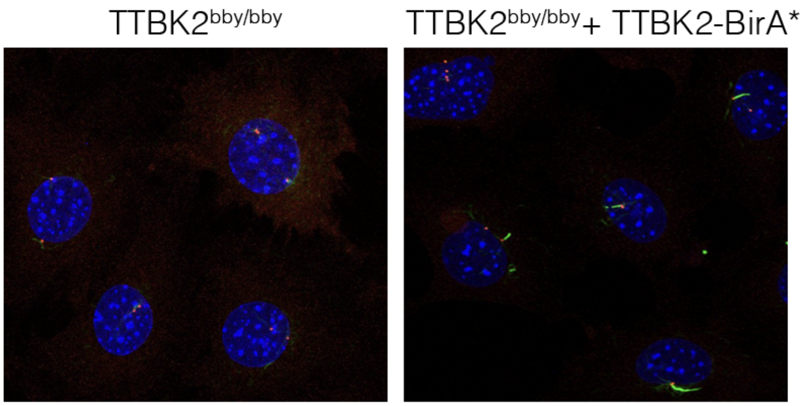


B

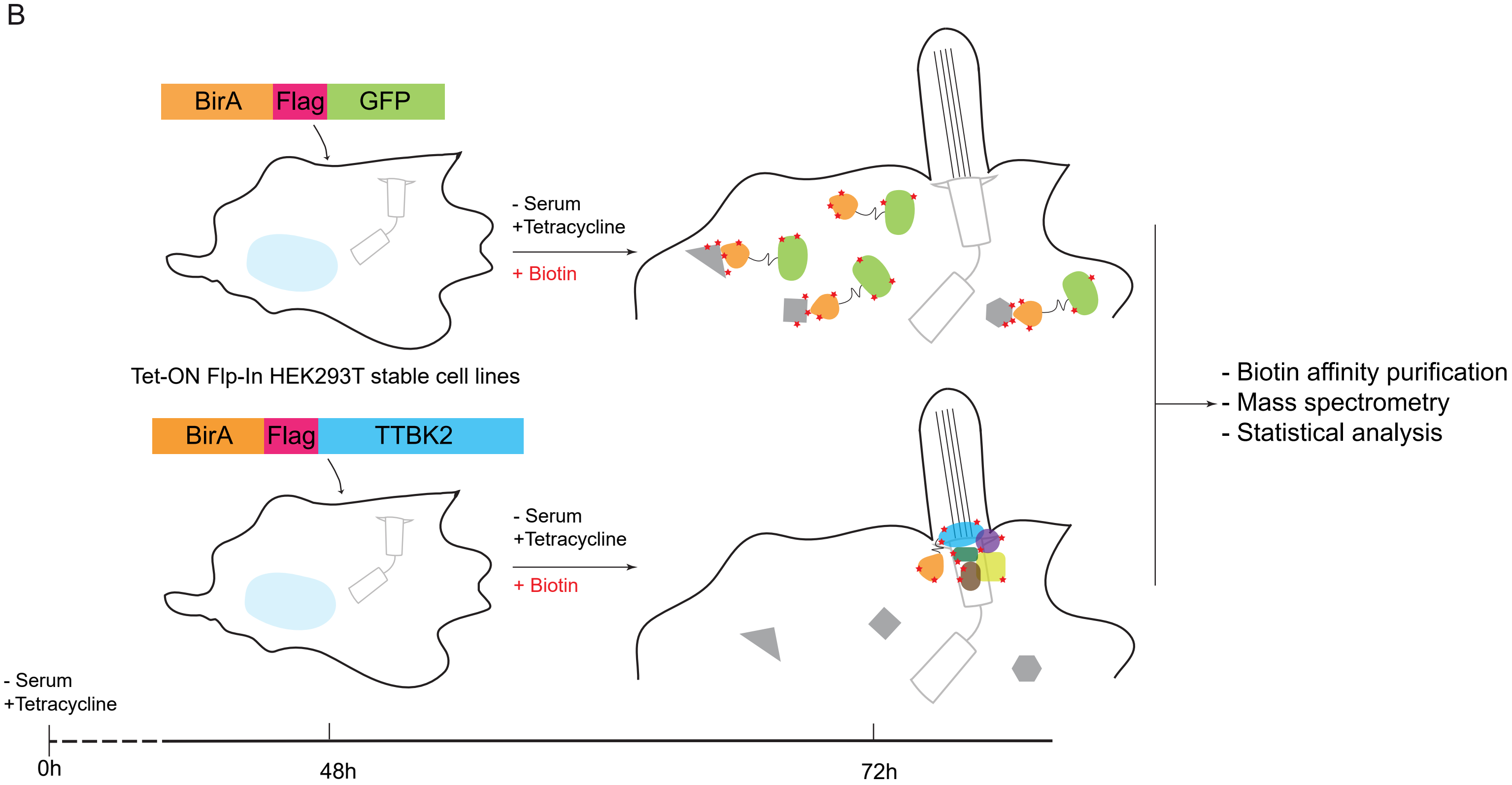


C

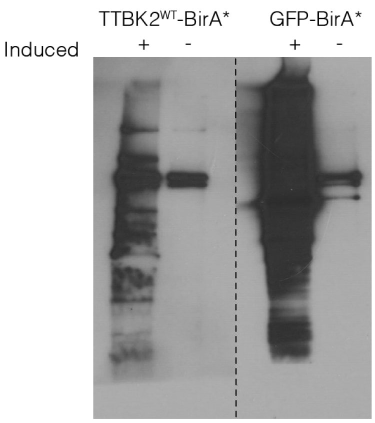

### Supplementary Figure S2

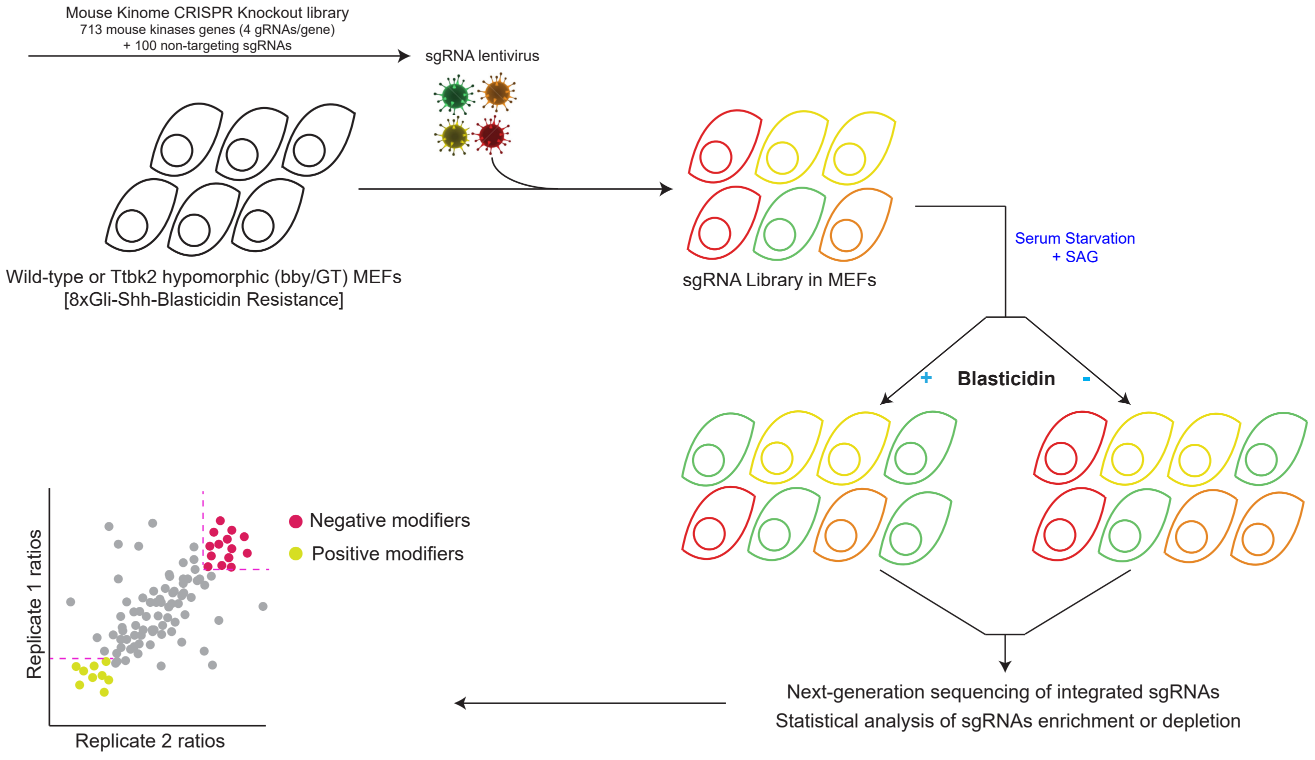

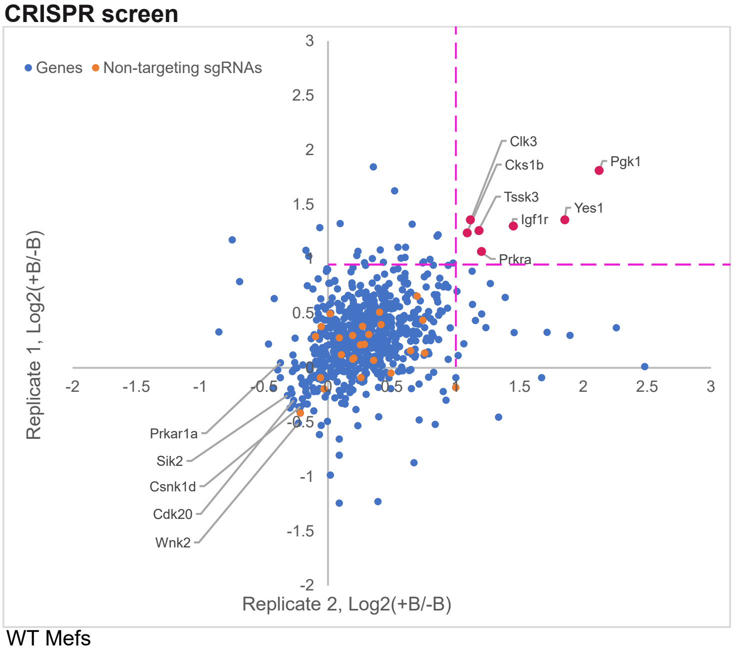

### Supplementary Figure S3

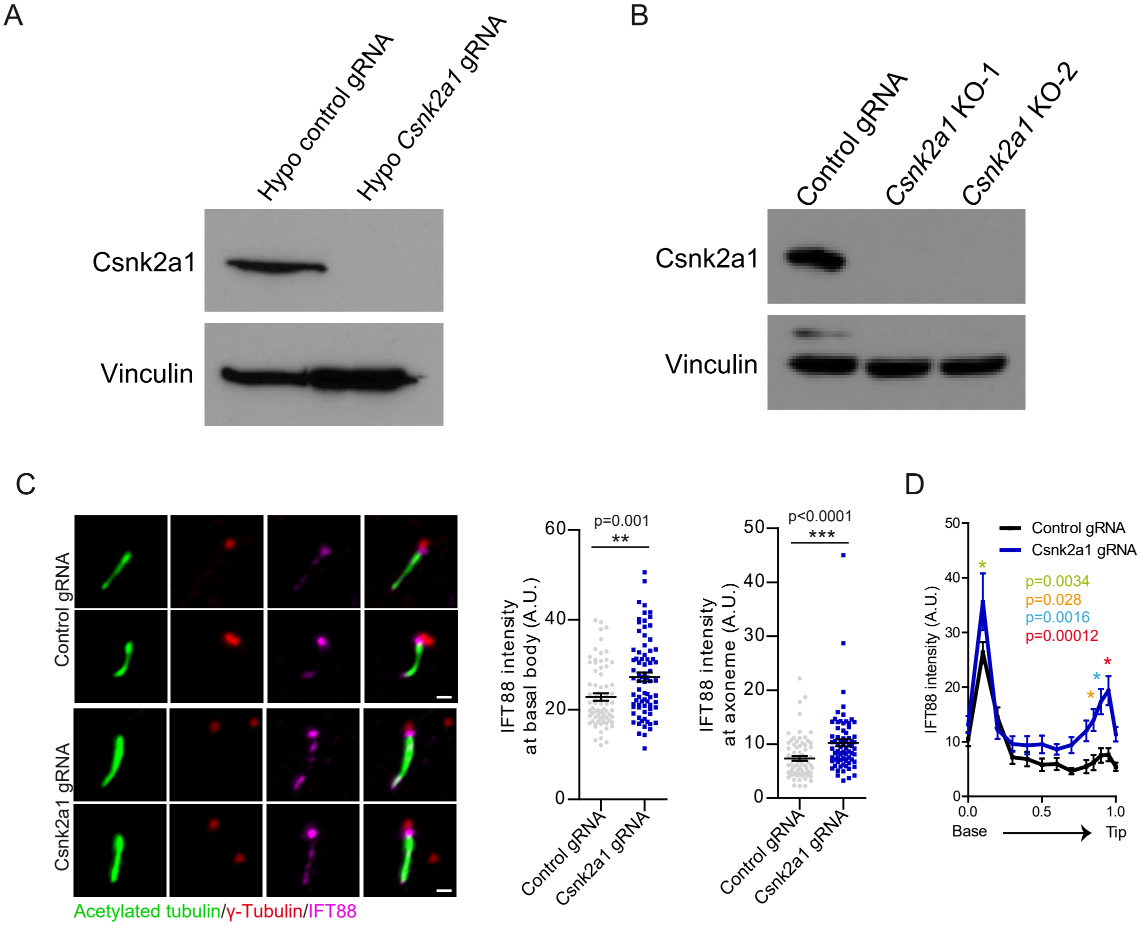

### Supplementary Figure S4

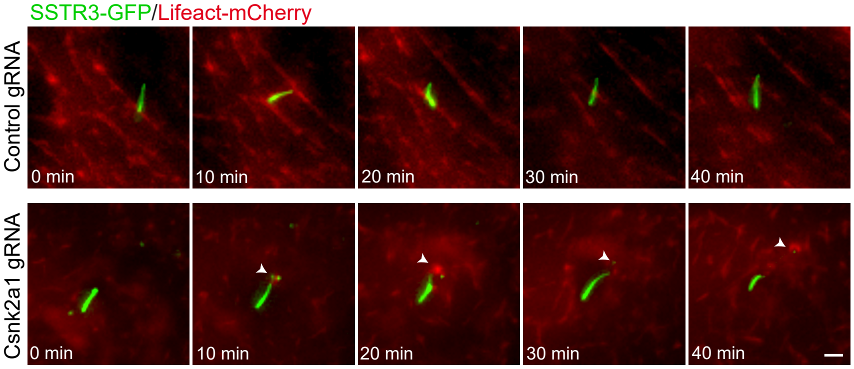

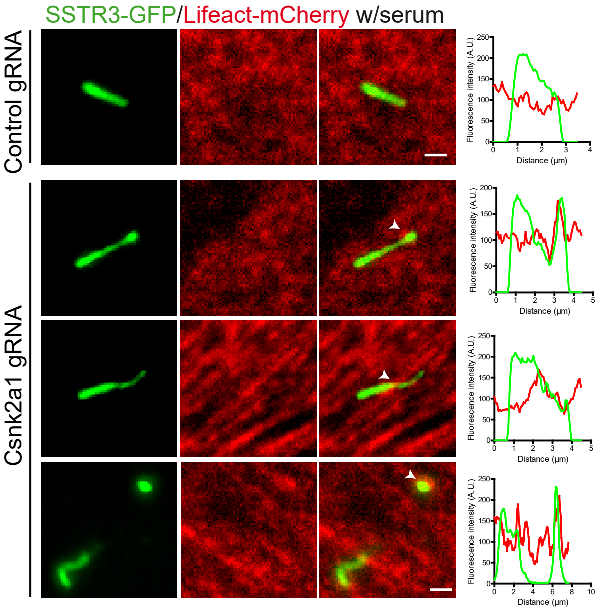
